# Cytosolic GLUCOSE-6-PHOSPHATE DEHYDROGENASE 5 is a key player in redox homeostasis during oxidative stress and in oxidative stress-triggered activation of the salicylic acid pathway

**DOI:** 10.64898/2026.05.01.722190

**Authors:** Lug Trémulot, Emmanuelle Issakidis-Bourguet, Katrien Van der Kelen, Bert De Rybel, Jean-Philippe Reichheld, Frank Van Breusegem, Graham Noctor, Amna Mhamdi

## Abstract

Glucose-6-phosphate dehydrogenase (G6PDH) catalyzes the first step of the oxidative pentose phosphate pathway, generating NADPH to sustain redox metabolism and signaling. However, whether individual G6PDH isoforms directly regulate oxidative stress signaling remains unclear. To determine the contribution of the different Arabidopsis G6PDH isoforms to oxidative stress signaling, we introduced single T-DNA mutants into the catalase-deficient *cat2* background, a genetic system in which intracellular H_2_O_2_ production activates salicylic acid (SA)-dependent cell death and defense pathways. Interestingly, impairment of cytosolic, but not chloroplastic G6PDH activity suppressed *cat2*-triggered phenotypes, with loss of *G6PD5* function fully abolishing lesion formation. The *cat2 g6pd5* double mutant phenocopied the SA biosynthesis-deficient mutant *cat2 sid2* and showed reversion of defense responses as well as metabolomic and transcriptomic profiles to the wild-type state. Strikingly, despite the suppression of SA-dependent lesions, loss of G6PD5 activity does not appear to reduce stress intensity. On the contrary, *cat2 g6pd5* plants exhibit increased glutathione synthesis and oxidation, elevated expression of oxidative stress marker genes, and enhanced accumulation of reactive nitrogen species relative to *cat2*. Protein-protein interaction analyses revealed that G6PD5 associates with several redox and defense-related proteins. In particular, we confirmed a physical interaction between G6PD5 and thioredoxin h5, a key component of redox-dependent SA signaling. However, analysis of *cat2 trxh5* and *cat2 npr1* lines indicated that this interaction alone cannot explain the G6PD5-dependent control of SA responses. Our work reveals that cytosolic G6PD5 integrates redox metabolism with immune signaling to control plant responses to oxidative stress.

## Introduction

In plant cells, maintaining redox homeostasis is a critical determinant of cellular integrity, enabling the coordination of metabolism with signaling under both optimal and stress conditions. Central to this network is reduced nicotinamide adenine dinucleotide phosphate (NADPH), a key reductant providing electrons for biosynthetic reactions and for the antioxidant systems that maintain cellular redox states (Foyer and Noctor, 2011; Lu et al., 2025). NADPH is generated through several metabolic routes (Noctor et al., 2015). These include photosynthetic electron transport in chloroplasts, which reduces ferredoxin that subsequently reduces NADP^+^ via ferredoxin-NADP^+^ reductase, and dehydrogenase reactions in primary metabolism, such as those catalyzed by cytosolic NADP-isocitrate dehydrogenase (cICDH) and NADP-malic enzyme (NADP-ME) (Foyer et al., 2011; Lu et al., 2025). A major contributor is the oxidative branch of the pentose phosphate pathway (OPPP), which generates two NADPH molecules per oxidized glucose-6-phosphate via the sequential actions of glucose-6-phosphate dehydrogenase (G6PDH) and 6-phosphogluconate dehydrogenase (6PGDH) (Kruger and von Schaewen, 2003).

NADPH plays a central role in cellular redox regulation by sustaining antioxidant and thiol-based redox systems. These include the ascorbate-glutathione pathway, where NADPH is used by monodehydroascorbate reductase (MDHAR) and glutathione reductase (GR) to regenerate ascorbate and glutathione, respectively (Foyer and Kunert, 2024; Noctor, 2025). NADPH also fuels thiol-dependent redox systems, involving thioredoxins and glutaredoxins which regulate protein redox state and peroxide detoxification (Dietz, 2011; Meyer et al., 2012; Geigenberger et al., 2017). In addition, NADPH serves as an electron donor for respiratory burst oxidase homologs (RBOHs) that generate reactive oxygen species (ROS) during development and responses to environment (Miller et al., 2010; Han et al., 2019; Kaya et al., 2019). Because ROS production and processing both depend on NADPH availability, metabolic control of NADPH generation represents a key regulatory node in oxidative stress signaling (Noctor et al., 2015).

To dissect how perturbations in intracellular hydrogen peroxide (H_2_O_2_) metabolism affect redox signaling, the catalase-deficient Arabidopsis mutant *cat2* has been widely used as a genetic model (Queval et al., 2007). Catalase 2 (CAT2) is a peroxisomal enzyme responsible for the disproportionation of H_2_O_2_ generated in the peroxisome, notably during photorespiration. Decreased catalase activity in *cat2* leads to increased engagement of other antioxidative pathways, particularly under conditions that allow significant photorespiratory flux such as ambient CO_2_ and standard light intensities (Queval et al., 2007; Mhamdi et al., 2010c). This conditionality makes *cat2* a powerful model to study how plants perceive and respond to oxidative signals linked to primary metabolism. Under photorespiratory conditions, *cat2* plants exhibit altered redox homeostasis, including increased glutathione oxidation and extensive transcriptional reprogramming. These changes activate salicylic acid (SA)-dependent defense responses, leading to expression of PATHOGENESIS-RELATED (*PR*) genes and the development of lesion-mimic phenotypes (Chaouch et al., 2010; Han et al., 2013). Genetic analyses combining *cat2* with mutations in redox-related pathways (e.g., *gr1*, *pad2*, *rbohF*) have underscored interactions between H_2_O_2_ metabolism, glutathione redox balance and SA signaling, highlighting the importance of NADPH-dependent processes in shaping oxidative stress (Mhamdi et al., 2010a; Chaouch et al., 2012; Han et al., 2013).

Several literature reports indicate that NADPH availability can modulate oxidative stress signaling. For example, combining *cat2* with mutations affecting NADPH-producing enzymes revealed that loss of cICDH exacerbates oxidative stress phenotypes, whereas loss of NADP-ME2 has little effect, suggesting distinct contributions of cytosolic NADPH sources (Mhamdi et al., 2010b; Li et al., 2013). In this context, G6PDHs, which catalyze the first and rate-limiting step of the OPPP, are likely major contributors to cytosolic NADPH supply. The Arabidopsis genome encodes six G6PDH isoforms, with G6PD1 to G6PD4 localized to plastids and G6PD5 and G6PD6 in the cytosol (Wakao and Benning, 2005). Modeling studies suggest that the OPPP may account for up to 65% of cytosolic NADPH production during oxidative stress (Tuzet et al., 2019). Consistent with this, G6PDHs have been implicated in various stress responses including salt tolerance, pathogen defense, and hormonal signaling involving salicylic acid and jasmonic acid (Stampfl et al., 2016; Hu et al., 2019; Yang et al., 2019; Née et al., 2023).

G6PDH is the first enzyme of the OPPP, a pathway that generates both NADPH and key carbon precursors such as ribose-5-phosphate and erythrose-4-phosphate to feed both primary and secondary metabolism (Kruger and von Schaewen, 2003). Importantly, G6PDH isoforms display distinct catalytic properties, and substitution of specific isoforms in other plant species can lead to marked physiological changes (Debnam et al., 2004; Wakao and Benning, 2005; Scharte et al., 2009; Scharte et al., 2023). However, despite these observations, the specific roles of individual G6PDH isoforms in oxidative stress signaling remain poorly characterized.

To investigate the role of G6PDH in oxidative stress signaling, we used the *cat2* system, in which oxidative stress triggers a higher demand for NADPH (Queval et al., 2007; Mhamdi et al., 2010c; Tuzet et al., 2019). Mutant lines for all six Arabidopsis *G6PD* genes were analyzed in conditions where increased oxidative stress triggers biotic stress phenotypes (Chaouch et al., 2010). Through this, we identify the cytosolic isoform G6PD5 as a key regulator of H_2_O_2_-triggered SA signaling. Loss of *G6PD5* function abolishes SA accumulation and lesion formation in *cat2*, causing phenotypes, transcriptomes, and metabolomes that copy those produced by directly impairing the SA-biosynthesis pathway. Further, G6PD5 associates with several redox and defense-related proteins, including thioredoxin h5 (TRXh5), and is required to promote redox homeostasis in oxidative stress conditions. Together, our findings highlight a central role for this protein in linking NADPH metabolism to redox signaling and immune responses during oxidative stress.

## Results

### The cytosolic G6PD5 plays a key role in transmitting oxidative stress signals

To investigate the functions of the different G6PDH isoforms, T-DNA mutants for each of the six G6PDH isoforms were obtained and characterized (Fig. 1A, Supplementary Figure S1). The expression of the corresponding gene was disrupted in all mutants, except for *g6pd2*. For this reason, this mutant was not used further. None of the mutations significantly impacted the transcript abundance of the other isoforms, excluding compensatory effects at this level. Measurement of G6PDH extractable activity in each of the lines revealed a decrease only in the cytosolic mutants, *g6pd5* and *g6pd6* (Fig. 1B). However, none of the mutations were associated with visible changes of the rosette phenotypes compared to Col-0 (Fig. 1C; top), suggesting functional redundancy between the different isoforms under standard conditions (long days (LD), 20 °C under an irradiance at leaf level of 200 µmol.m^-2^.s^-1^).

**Figure 1.**
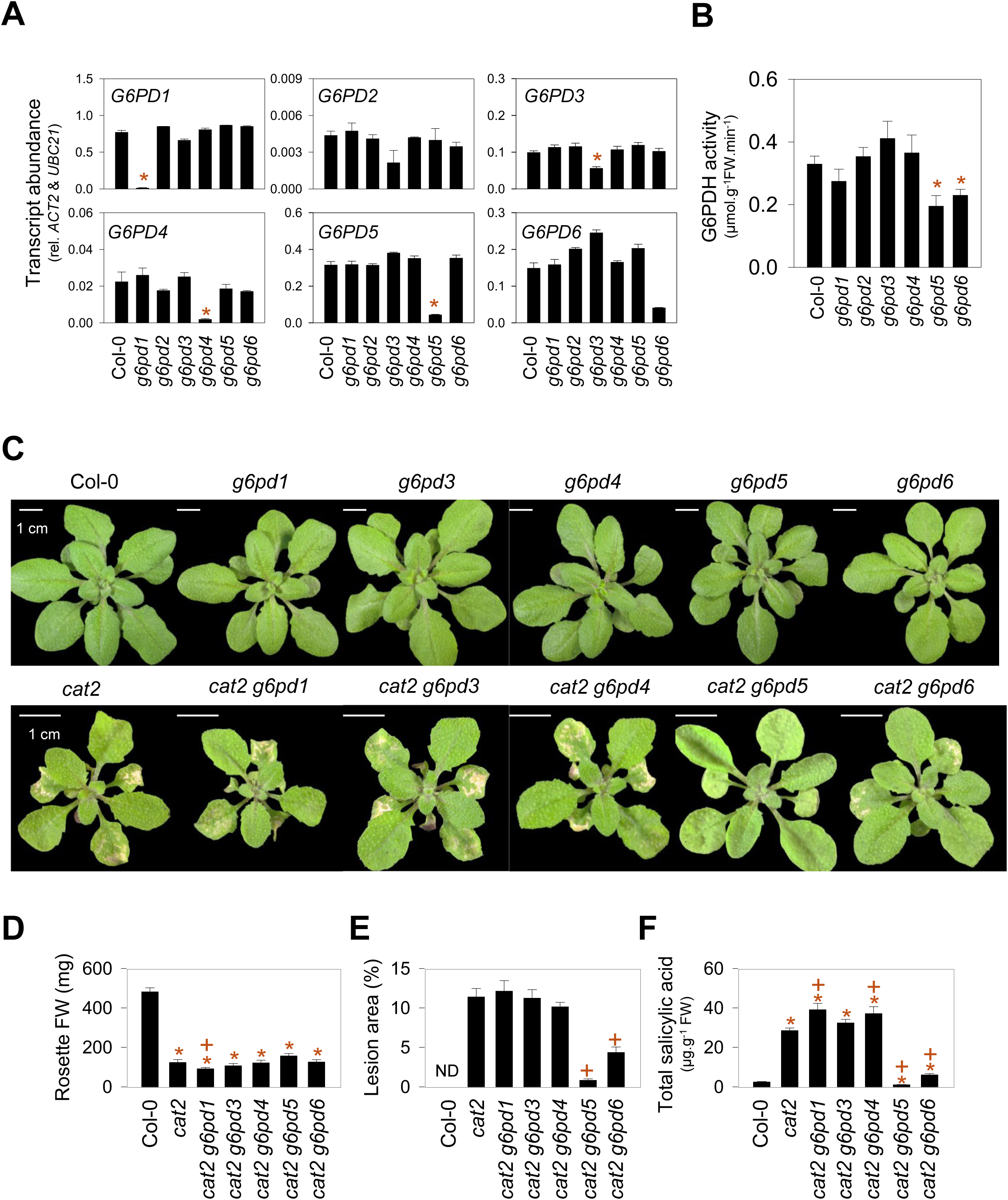
Specific loss of *G6PD5* function blocks oxidative stress-induced immune responses in Arabidopsis. A. Analysis of *G6PD* transcript abundance in respective single mutants. Transcript abundance was determined by RT-qPCR and normalized to *ACT2* and *UBC21* reference genes. B. Total extractable G6PDH activity in the different single mutants. In A and B, data are means ± SE of three biological replicates. C. Representative photographs of Arabidopsis single and double mutants. D. Fresh weight measurements (n=12 plants per genotype). E. Lesion area quantified as a percentage of total rosette area. F. Total salicylic acid (SA) levels measured in 6 biological replicates. In D, E and F, data are means ± SE of biological replicates as indicated above. Plants were grown in long days for three weeks. Asterisks (*) indicate significant differences relative to Col-0, and plus signs (+) indicate significant differences relative to *cat2* (Student’s *t*-test, *p* < 0.05).

Our aim in initiating this project was to evaluate the role of the different G6PDH isoforms in oxidative stress conditions. To this end, the different single mutations were introduced into the genetic background *cat2*, where oxidative stress intensity can be sustained and easily controlled by external growth conditions (e.g., CO_2_ levels and light intensity). As previously reported, *cat2* presented visible lesions on the leaves, as well as a dwarfed rosette (Fig. 1C-E) when grown in standard conditions (LD, in air from seeds, light intensity of 200 µmol.m^-2^.s^-1^ at leaf level). The leaf phenotypes of *cat2 g6pd1*, *cat2 g6pd3*, and *cat2 g6pd4* were similar to those observed in the *cat2* single mutant (Fig.1C-F). However, the *cat2 g6pd6* mutants showed fewer lesions compared to *cat2* and decreased accumulation of SA (Fig. 1C, bottom panel: Fig.1-F). Most strikingly, no lesions were apparent on the leaves of *cat2 g6pd5* (Fig. 1C; bottom panel and Fig. 1E).

The absence of lesions in *cat2 g6pd5* was associated with complete abolition of the *cat2*-triggered accumulation of leaf SA (Fig. 1E). To confirm that the suppression of the *cat2*-triggered lesions and SA contents were specifically due to loss of *G6PD5* function, the mutant was complemented with a construct expressing G6PD5 fused to GFP under the control of the 35S promoter. Two independent complemented lines, *cat2 g6pd5 G6PD5-1* and *cat2 g6pd5 G6PD5-2*, both showed fully restored lesion phenotypes similar to *cat2* (Fig. 2A). Complementation was accompanied by a marked increase in extractable G6PDH activity relative to *cat2 g6pd5*, *cat2*, and Col-0 (Fig. 2B). It is also noteworthy that *cat2* itself displayed higher G6PDH activity than Col-0, consistent with enhanced engagement of the OPPP in oxidative stress conditions (Fig. 2B). In the complemented lines, the restoration of the lesion phenotype was associated with SA accumulation and increased expression of the SA synthesis gene *ICS1* and the SA marker gene *PR1* (Fig. 2C). Strikingly, extractable G6PDH activity positively correlated with total SA levels, further supporting a functional link between cytosolic G6PD5 activity and SA signaling during oxidative stress (Fig. 2B-C). To further verify the role of *G6PD5*, an allelic loss-of-function mutant, *cat2 g6pd5-2,* was analyzed (Supplementary Figure S2 A-B). The *cat2 g6pd5-2* line did not develop visible lesions and exhibited lower *PR1* expression and SA accumulation compared to *cat2* (Supplementary Figure S2 C-D). These results identify cytosolic G6PDH activity, and G6PD5 in particular, as a key component determining SA-associated responses in the oxidative stress background, *cat2*.

**Figure 2.**
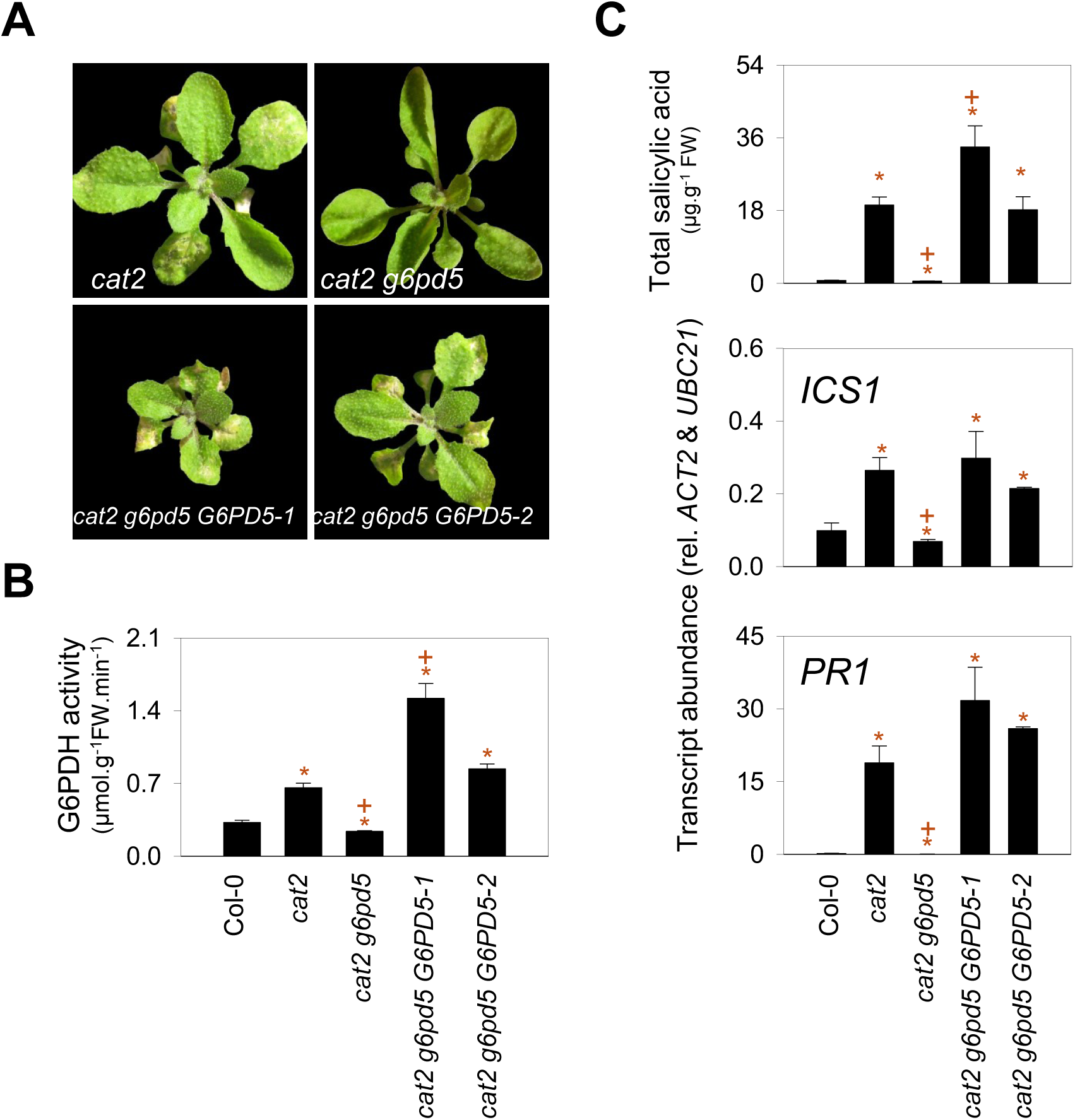
Complementation of the *cat2 g6pd5* mutants restores lesion formation and SA signaling. A. Phenotypes of *cat2 g6pd5* complemented lines. B. Total extractable G6PDH activity in the different genotypes. C. Salicylic acid contents and *ICS1* and *PR1* transcript abundance. Transcript abundance was determined by RT-qPCR and normalized to *ACT2* and *UBC21* reference genes. Plants were grown in long days for three weeks. Data are means ± SE of three biological replicates. Asterisks (*) indicate significant differences relative to Col-0, and plus signs (+) indicate significant differences relative to *cat2* (Student’s *t*-test, *p* < 0.05).

### Loss of *G6PD5* mimics loss of *SID2* and abolishes oxidative stress-triggered responses

The phenotypes observed in the *cat2 g6pd5* lines suggest that oxidative stress no longer entrains SA accumulation when *G6PD5* expression is impaired. We therefore conducted a direct comparison with the previously reported *cat2 sid2* double mutant (Chaouch et al., 2010). The *sid2* mutation in ISOCHORISMATE SYNTHASE 1 causes a defect in SA synthesis: as a result, *cat2 sid2* does not show lesions in oxidative stress conditions (Chaouch et al., 2010).

Although similar rosette fresh weights were measured in *cat2*, *cat2 sid2* and *cat2 g6pd5* (Supplementary Figure S3A), both double mutants failed to develop lesions (Fig. 3A) or accumulate SA (Fig. 3B). Indeed, SA remained at levels below those of Col-0 in both double mutants (Fig. 3B, Supplementary Figure S3B). These decreased SA levels were accompanied by lower accumulation of the defense-related metabolites scopoletin and camalexin (Fig. 3B; Supplementary Figure S3B). Consistent with these changes, both *cat2 sid2* and *cat2 g6pd5* exhibited increased growth of *Pseudomonas syringae* DC3000 compared to *cat2* and Col-0, indicating enhanced susceptibility to infection (Supplementary Figure S3C). Together, these results show that loss of G6PD5 phenocopies the SA-deficient *sid2* mutant and affects lesion formation, SA accumulation, defense-related metabolites, and pathogen responses.

**Figure 3.**
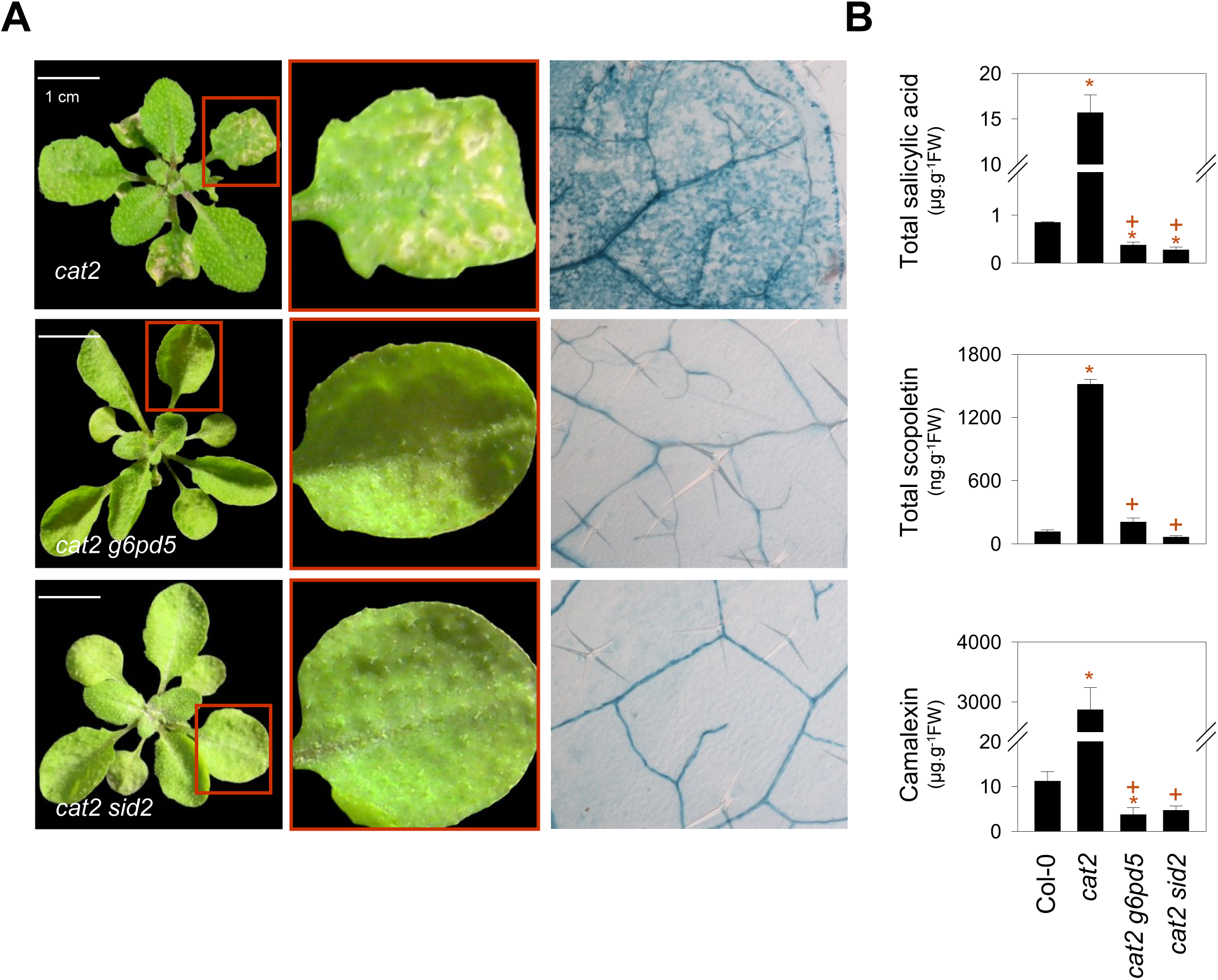
The *g6pd5* mutation mimics the *sid2* mutation. A. Photographs of plants growing in long days for three weeks and trypan blue staining of selected leaves. B. Defense metabolites in *cat2 g6pd5* and *cat2 sid2* mutants. Data are means ± SE of three biological replicates. Asterisks (*) indicate significant differences relative to Col-0, and plus signs (+) indicate significant differences relative to *cat2* (Student’s *t*-test, *p* < 0.05).

### The *g6pd5* mutation reverts the metabolic and molecular features triggered by *cat2* in a similar manner to *sid2*

To gain insight into the role of G6PD5, untargeted metabolomic and transcriptomic analyses were performed in Col-0, *cat2*, and *cat2 g6pd5*. To further evaluate relationships between G6PD5 and the SA pathway, the *cat2 sid2* mutant was included for comparison with *cat2 g6pd5*.

We previously reported that oxidative stress in *cat2* drives specific changes in primary and secondary metabolism, including increased accumulation of several amino acids and defense-related hormones (Noctor et al., 2015; Lelarge-Trouverie et al., 2023). By contrast, the metabolic profile of *cat2 g6pd5*, and similarly *cat2 sid2*, more closely resembled that of Col-0 than *cat2*; ie, a large part of the *cat2*-driven changes was annulled (Fig. 4A; Supplementary Table S2). Notably, gluconic acid and nicotine were the two metabolites showing the strongest and most significant decrease in *cat2 g6pd5* relative to *cat2*, and their levels were comparable in *cat2 g6pd5* and *cat2 sid2* (Fig. 4B). On the other hand, *myo*-inositol, one of the few metabolites with decreased contents in *cat2*, was more accumulated in the double mutants, although to a lesser extent than in Col-0 (Fig. 4B). Thus, the introduction of *g6pd5* into the *cat2* background partly annuls oxidative stress-triggered metabolic changes, with high similarity to effects produced by the *sid2* allele.

**Figure 4.**
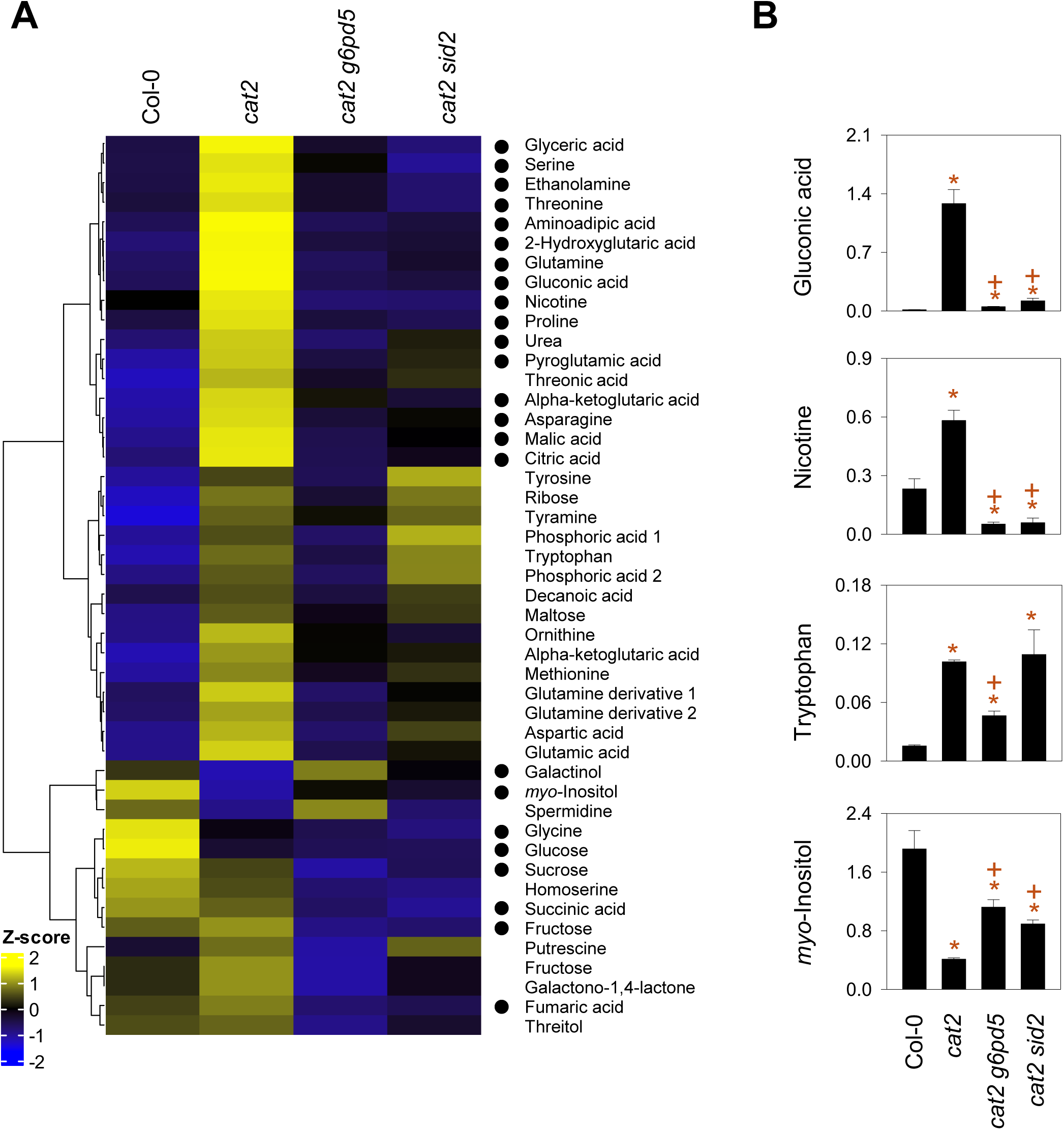
Metabolome profiles in *cat2 g6pd5* and *cat2 sid2.* A. Heatmap of metabolites quantified by GC-MS in Col-0, *cat2*, *cat2 sid2* and *cat2 g6pd5* (Data are shown as z-scores). Metabolites displayed (46 metabolites) were statistically different between *cat2 g6pd5* and *cat2*. The black dot aside the metabolite name indicates a statistically different abundance between *cat2 sid2* and *cat2*. B. Examples of metabolites that were differentially abundant in *cat2 g6pd5* compared to *cat2*. Mean peak intensities of selected metabolites in the four genotypes. Data show raw values for characteristic MS fragments corrected for fresh weight and internal standards. For easier display, the intensities of each metabolite were multiplied by 1000 in B. Data are means ± SE of three biological replicates. Asterisks (*) indicate significant differences relative to Col-0, and plus signs (+) indicate significant differences relative to *cat2* (Student’s *t*-test, *p* < 0.05).

The effects of the *cat2* single mutation on transcriptome profiles have previously been well documented (Queval et al., 2012; Willems et al., 2016; Lelarge-Trouverie et al., 2023; Xu et al., 2025), whereas the *cat2 sid2* transcriptome has not been reported previously. Here, we analyzed transcriptomic changes in Col-0, *cat2*, *cat2 g6pd5*, and *cat2 sid2*. Principal component analysis and differential gene expression highlighted a clear separation between *cat2* and other genotypes including *cat2 g6pd5* and *cat2 sid2*, which clustered closely together (Supplementary Figure S4; Supplementary Table S3).

Analysis of the 1814 differentially expressed genes (DEGs) identified in the Col-0 versus *cat2* comparison revealed that *cat2 g6pd5* resembled Col-0 and exhibited expression profiles largely opposite to those of *cat2*, a trend that was also observed in *cat2 sid2* (Fig. 5A). The top 30 genes showing the highest fold change between Col-0 and *cat2* were strongly down-regulated in both *cat2 g6pd5* and *cat2 sid2*, with the oxidative stress marker, *GSTU24*, as a notable exception (Fig. 5B), once more reflecting the high similarity between these two genotypes.

**Figure 5.**
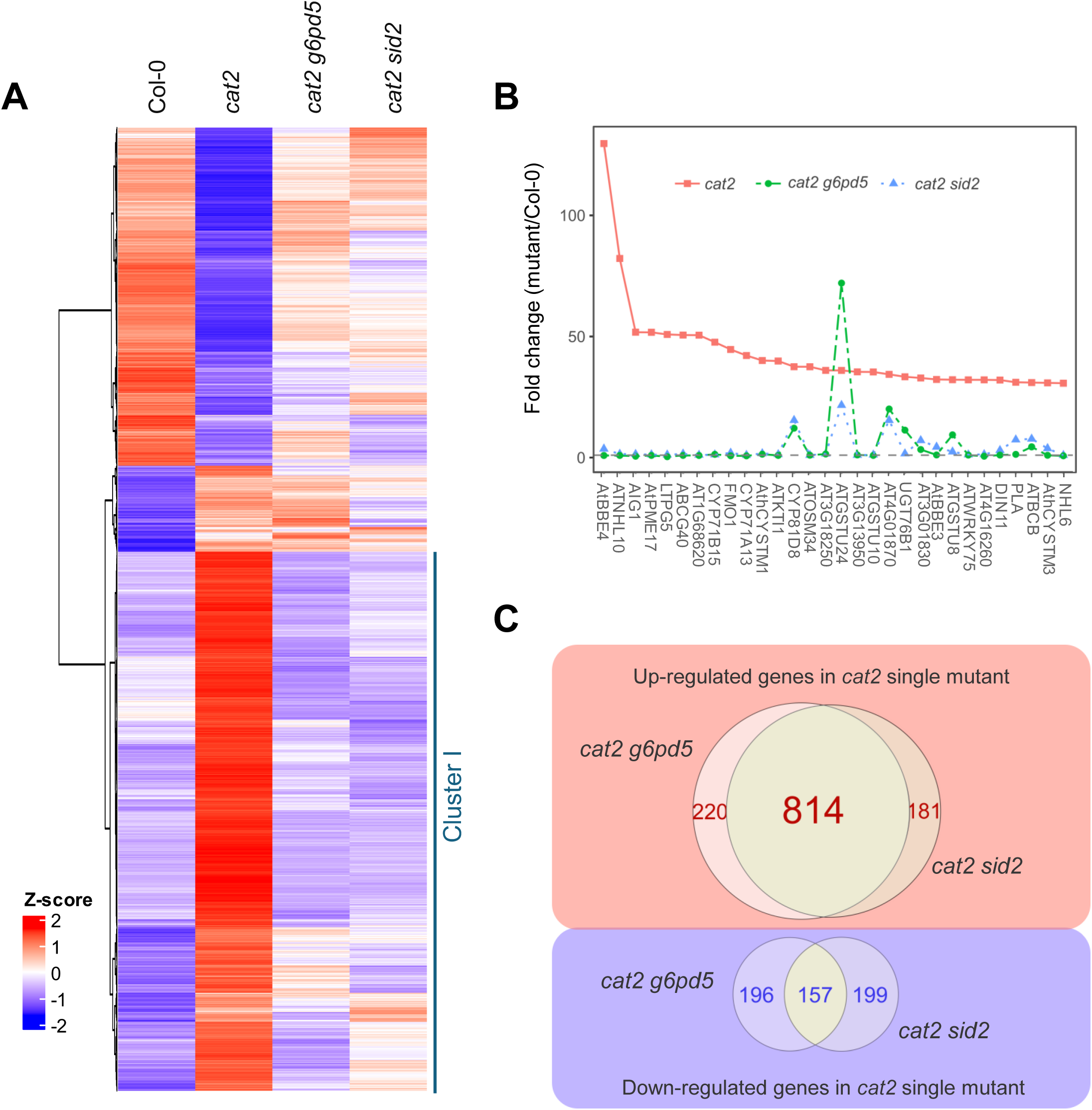
Transcriptome analysis of *cat2 g6pd5* and *cat2 sid2*. A. Heatmap showing gene expression profiles Col-0, *cat2*, *cat2 g6pd5 cat2 sid2*. Displayed genes are the 1814 *cat2* vs Col-0 DEGs in the microarray experiments, shown as Z-scores. The dendrogram corresponds to Pearson correlation with Ward.D2 distance. B. Effect of additional mutations (*g6pd5* and *sid2*) on the expression of the top 30 genes induced in the *cat2* background. The fold change is determined using the mean of intensity with Col-0 as reference. C. Venn diagrams showing overlap of DEGs in *cat2 g6pd5* and *cat2 sid2*. contrasts comparing *cat2* with *cat2 sid2* or *cat2 g6pd5*. The figures indicate the number of DEGs within each specific set. For example, 220 genes were differentially expressed between *cat2 g6pd5* and *cat2* that did not also show differential expression between *cat2* and *cat2 sid2* while 814 genes were found to be differentially expressed in the two comparisons.

Consistent with these observations, direct comparisons of *cat2 g6pd5* versus *cat2* and *cat2 sid2* versus *cat2* revealed a strikingly high degree of overlap between the two double mutants. Notably, more than three-quarters of the up-regulated DEGs were shared between *cat2 g6pd5* and *cat2 sid2* (Fig. 5C), indicating that both mutations trigger highly similar transcriptional reprogramming in the *cat2* background. Gene Ontology enrichment analysis of DEGs up-regulated in *cat2* and down-regulated in Col-0 (Cluster I, Fig. 5A) revealed significant enrichment for biotic stress-related processes (Supplementary Figure S5B). Among the most strongly affected genes in the *cat2 g6pd5* versus *cat2* comparison, *HSP17.6C* (*SMALL HEAT-SHOCK PROTEIN*) and *XTR8* (*XYLOGLUCAN ENDO-TRANSGLYCOSYLASE-RELATED 8*) were among the most highly up-regulated, whereas *PR1* and *AtBBE4* (*FAD-BINDING BERBERINE FAMILY PROTEIN*) were among the most strongly down-regulated (Supplementary Figure S5B). Interestingly, *HSP17.6C* showed a specific induction in the *cat2 g6pd5* mutants, whereas the other genes displayed similar expression patterns in *cat2 g6pd5* and *cat2 sid2* relative to *cat2*.

Together, these analyses show that loss of *G6PD5* function closely mirrors the *sid2* mutation and opposes the *cat2*-induced responses, thereby largely reverting both metabolic and transcriptional profiles towards the wild-type levels.

### Redox homeostasis is substantially altered in *cat2 g6pd5*

Because G6PD5 contributes to NADPH production, loss of its activity might be predicted to influence cellular redox homeostasis under conditions of oxidative stress. This hypothesis receives some support from the above-described morphological, molecular, and metabolic shifts. Analysis of key redox factors provided more direct evidence. The leaf NADPH/NADP⁺ ratio was altered in *cat2 g6pd5*, displaying values intermediate between those observed in Col-0 and *cat2* (Supplementary Figure S6). By contrast, neither NAD(H) pools nor the ascorbate pool size and redox state were affected significantly in *cat2 g6pd5* or *cat2* (Supplementary Figure S6). By striking contrast, glutathione metabolism was markedly affected. Already higher and more oxidized as a result of the oxidative stress in *cat2*, glutathione accumulated to very high levels in the *cat2 g6pd5* double mutant, and was very highly oxidized (Fig. 6A). The glutathione precursors cysteine (Cys) and γ-glutamyl-cysteine (γ-EC) also accumulated in *cat2 g6pd5* and *cat2*, although the differences between *cat2* and *cat2 g6pd5* were smaller than those observed for glutathione (Fig. 6B). In addition, the glutathione degradation product cysteinyl-glycine, Cys-Gly, accumulated to higher levels in *cat2 g6pd5* than in either *cat2* or Col-0. Together, these data suggest that in conditions of oxidative stress, G6PD5 is required to limit glutathione oxidation and accumulation, together with the attendant changes in synthesis and turnover.

**Figure 6.**
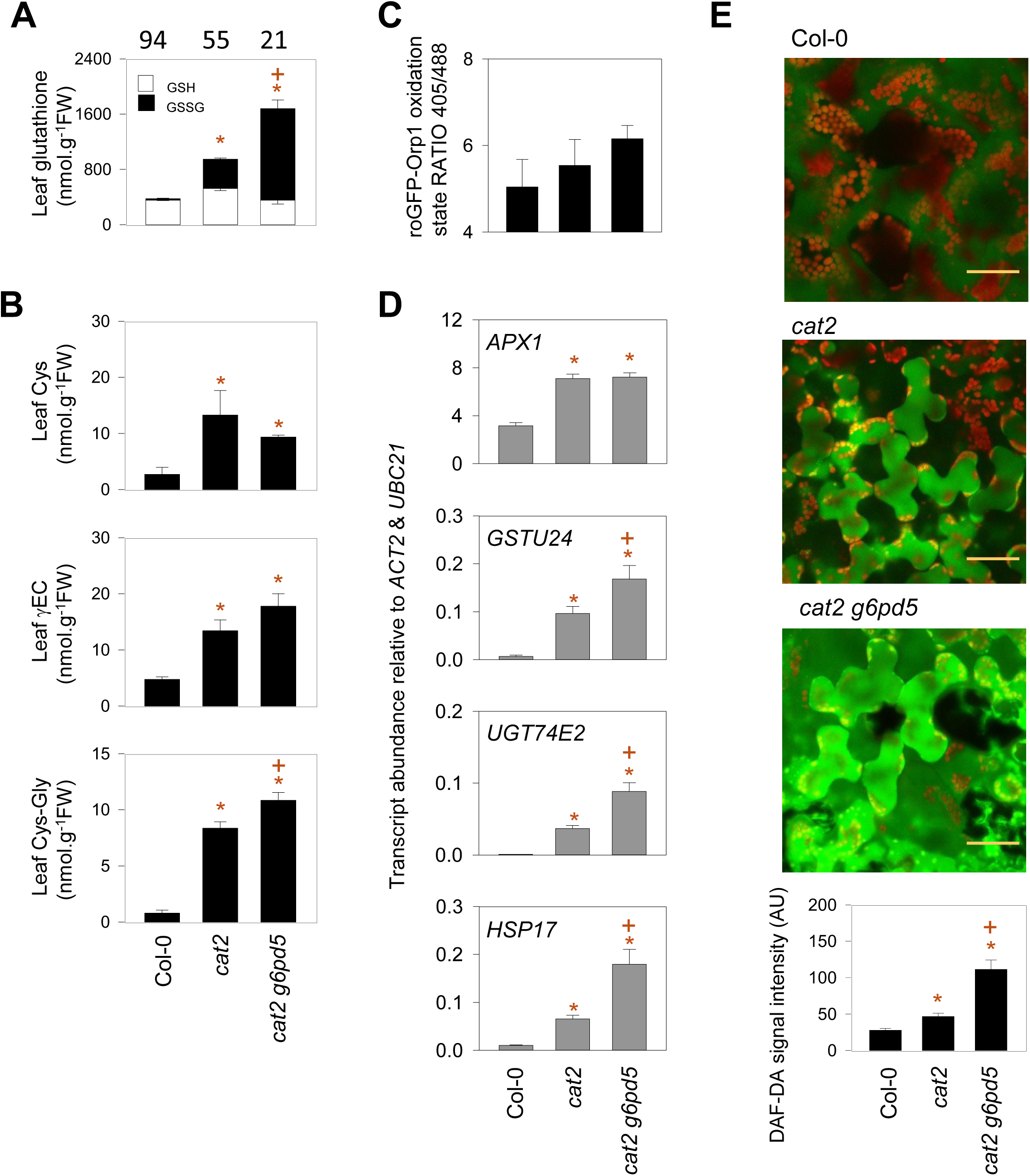
Loss of G6PD5 alters redox homeostasis and signaling. A. Glutathione pool and redox state in Col-0, *cat2*, and *cat2 g6pd5.* Reduced glutathione and oxidized glutathione are indicated by white boxes and black boxes, respectively. Numbers above bars indicate the percentage of glutathione present in its reduced form. B. Contents of glutathione biosynthesis precursors and degradation products. C. Degree of oxidation of the genetically encoded H₂O₂ sensor roGFP2-Orp1, expressed as the 405/488 nm fluorescence ratio, measured in leaf discs using a CLARIOstar plate reader. Data are means ± SE of thirty biological replicates (different leaf disks). Higher ratios indicate increased probe oxidation. D. Transcript abundance of oxidative stress marker genes *APX1*, *GSTU24*, *UGT74E2*, and *HSP17*, determined by RT-qPCR and normalized to *ACT2* and *UBC21*. D. Accumulation of reactive nitrogen species assessed by DAF-2DA fluorescence in leaf tissue. Representative confocal images are shown, with chlorophyll autofluorescence displayed in red. Scale bars = 50 µm. Quantification of DAF-2DA fluorescence is shown below the photographs. Data are means ± SE of fourteen biological replicates. For easier display, fluorescence values were divided by a factor of 100,000. In A, B and D, data are means ± SE of three biological replicates. Asterisks (*) indicate significant differences relative to Col-0, and plus signs (+) indicate significant differences relative to *cat2* (Student’s *t*-test, *p* < 0.05).

Given the cytosolic localization of G6PD5, we used the cytosolic H_2_O_2_-sensitive roGFP-Orp1 probe (Nietzel et al., 2019) to assess the concentration of H_2_O_2_ in the different genotypes (Fig. 6C). Despite the clear redox effects apparent from glutathione measurements, this analysis revealed no obvious marked increase in roGFP-Orp1 oxidation in *cat2* or *cat2 g6pd5* (Fig. 6C). It should be noted that this observation is consistent with previous measurements of extractable H_2_O_2_ levels in *cat2* grown in similar conditions, which were not greatly increased relative to the wild-type (Noctor et al., 2015). Despite this, oxidative stress marker genes are consistent with the glutathione data in showing oxidative stress in *cat2* (Mhamdi et al., 2017; Noctor et al., 2023) and were therefore used to assess the effects of the *g6pd5* mutation (Fig. 6D). While *APX1* transcripts were similar in *cat2* and *cat2 g6pd5*, another three marker genes (*GSTU24*, *UGT74E2*, and *HSP17*) accumulated to significantly higher levels in the double mutant (Fig. 6D). In the case of *GSTU24*, this effect is consistent with the RNAseq data described above (Fig. 5B). In parallel, the fluorescence of the reactive nitrogen species (RNS)-sensitive probe DAF-DA was increased in *cat2 g6pd5* relative to both *cat2* and Col-0 (Fig. 6E). Collectively, these results indicate that loss of *G6PD5* function significantly affects redox regulation and that abolition of SA signaling in *cat2 g6pd5* cannot be explained by decreased oxidative stress intensity compared to *cat2*.

### G6PD5 interacts with proteins related to biotic stress response and redox homeostasis

Given the apparent importance of G6PD5 in transmitting oxidative stress signals, we investigated whether this function might involve interactions with other proteins. A pull-down strategy based on immunoprecipitation followed by mass spectrometry (IP-MS/MS, Wendrich et al., 2017) was performed using GFP-tagged G6PD5 expressed in a complemented *cat2 g6pd5* line (*cat2 g6pd5* G6PD5-2). To restrict the analysis to robust candidate interactors, stringent selection criteria were applied, including an enrichment ratio greater than 50 and a significance threshold below 0.01 relative to the *cat2* control. This filtering resulted in the identification of 90 putative interacting proteins (Supplementary Table S4; Table 1). Functional enrichment analysis of this dataset revealed a significant overrepresentation of terms related to biotic stress responses and heat responses (Fig. 7B). Annotation-based grouping further indicated that these proteins participate in diverse biological processes (Fig. 7C), with a notable enrichment of redox- and defense-related candidates, including GSH1, GSNOR, TRXh5, PR1, and PR5. Of these, PR1, PR5, and TRXh5 showed the highest enrichment values (Table 1).

**Figure 7.**
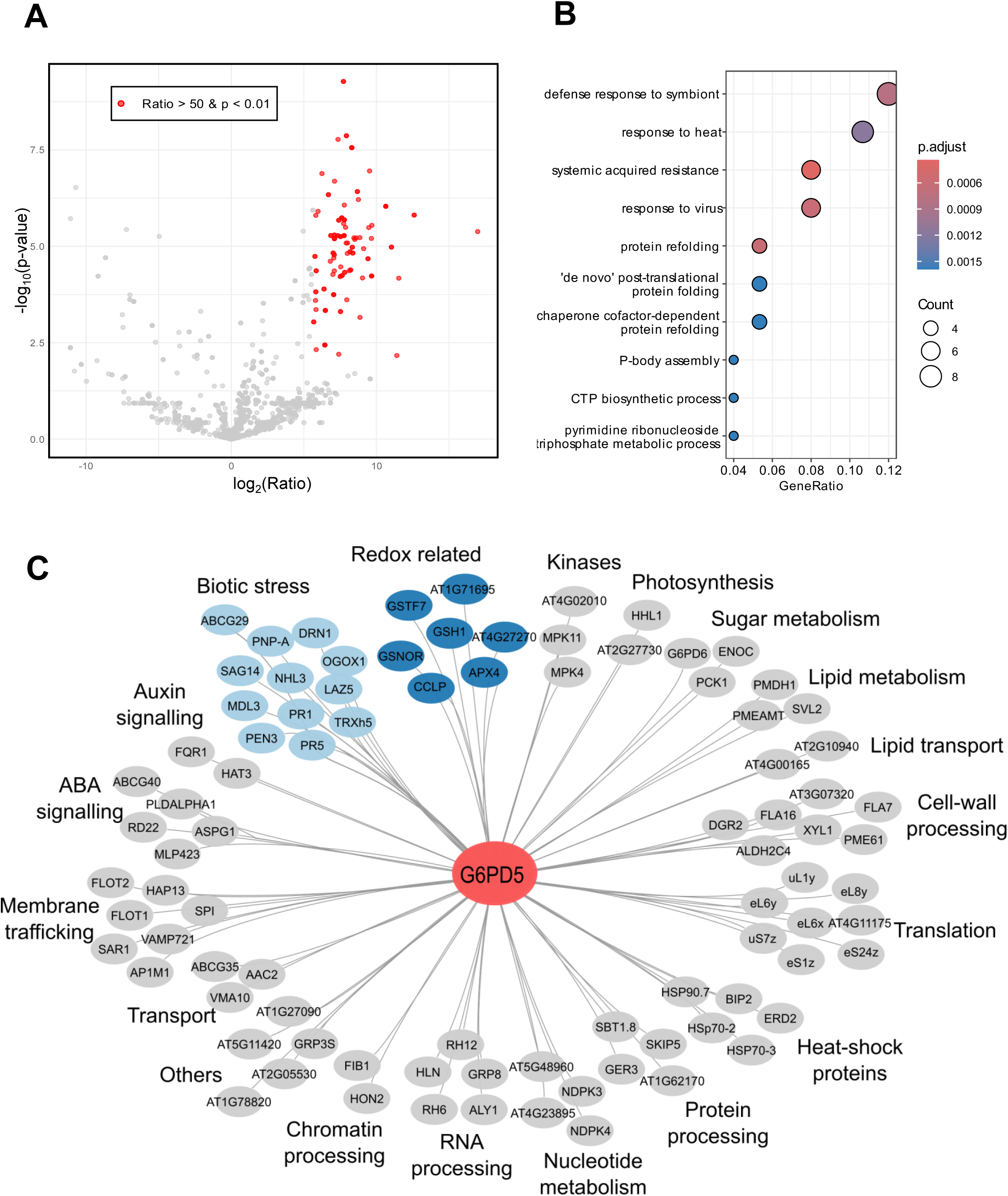
Potential G6PD5 interacting proteins identified by IP-MS. Volcano plot of proteins putatively interacting directly or indirectly with G6PD5 identified by a pulldown-MS/MS strategy. Candidates with an enrichment ratio higher than 50 and a *p*-value lower than 0.01 are coloured in red. Data corresponds to peptides identified by MS/MS assigned to proteins and enrichment refers to the comparison between *cat2 g6pd5* G6PD5-GFP and *cat2* (control). A. Gene ontology (GO) enrichment for the list of 90 candidate proteins interacting with G6PD5. Enrichments are displayed for the top 10 GO biological processes. Dot plots represent GO terms enriched according to GeneRatio, which gives the ratio between the number of genes coding for proteins responsible for enrichment and the total number of genes annotated for the GO term. Dot size corresponds to the number of genes responsible for the enrichment, and dot colors to the *p*-values for the hypergeometric distribution. C. Cytoscape network of potential interacting proteins. Proteins are clustered and colored according to biological function based on annotation and the literature. Each black edge corresponds to a probable direct or indirect interaction.

**Table 1.**
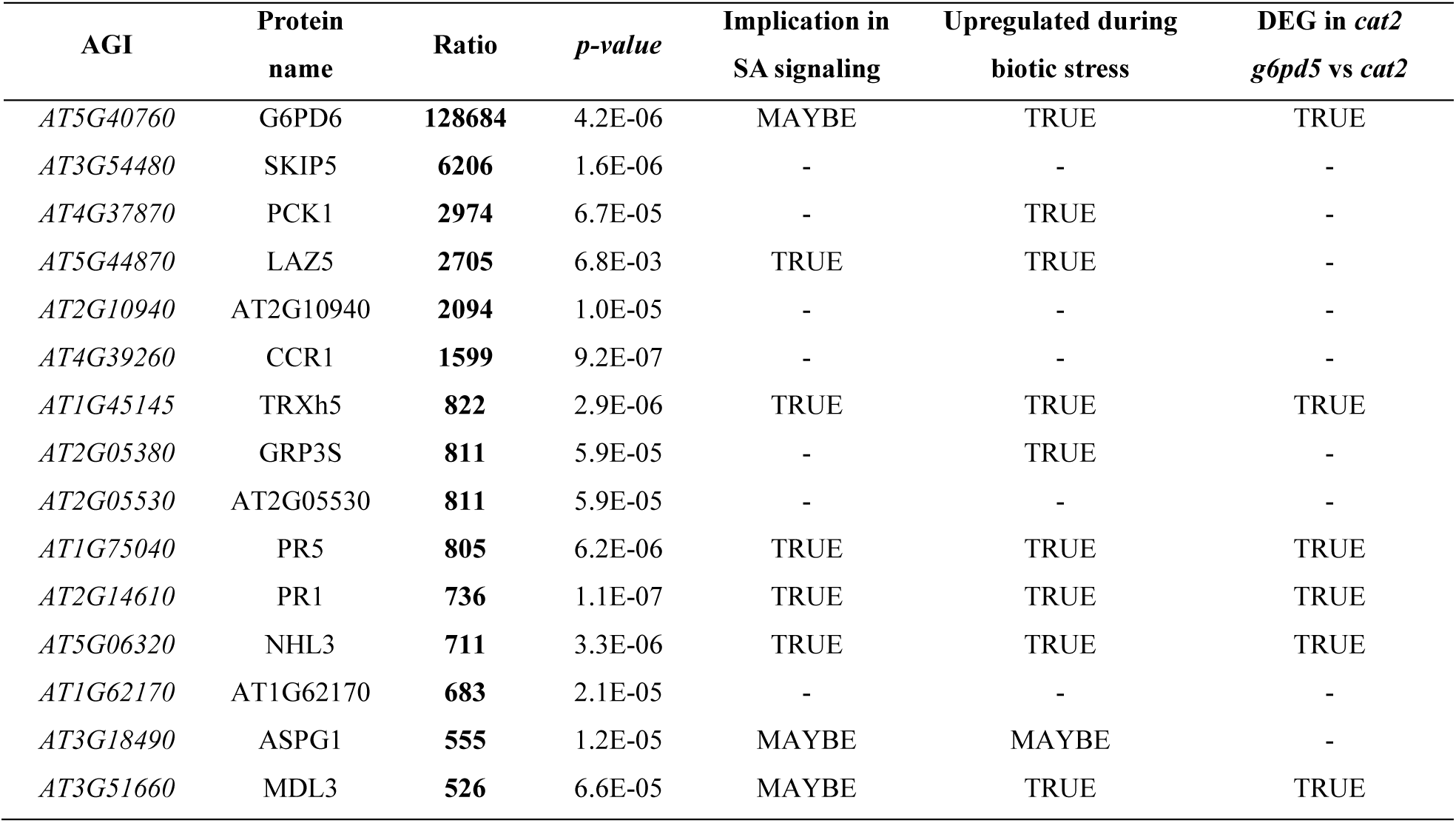
Top enriched proteins in *cat2 g6pd5 G6PD5-GFP* samples. Proteins showing strong enrichment over the control (ratio > 500; *p* < 0.01) are listed, highlighting the most highly enriched interactor candidates. Functional links to salicylic acid signaling and biotic stress responses were inferred using ThaleMine and ePlant. Differentially expressed genes in the *cat2 g6pd5* versus *cat2* comparison are indicated.

Among the identified interactors, TRXh5 was selected for further analysis because of its reported role in SA signaling and its NADPH-dependent redox activity via NADPH-thioredoxin reductase (NTR; Laloi et al., 2004; Tada et al., 2008; Geigenberger et al., 2017). The interaction between G6PD5 and TRXh5 was tested by transient co-expression in *Nicotiana benthamiana*, followed by immunoblot analysis. This approach confirmed the physical interaction between G6PD5 and TRXh5 (Fig. 8A). In parallel, *TRXh5* transcript and protein abundance were quantified in Col-0, *cat2*, and *cat2 g6pd5*. Both *TRXh5* transcript and TRXh5 protein levels were increased in *cat2* but not in the double mutant *cat2 g6pd5* (Fig. 8B and C, Supplementary Figure S7).

**Figure 8.**
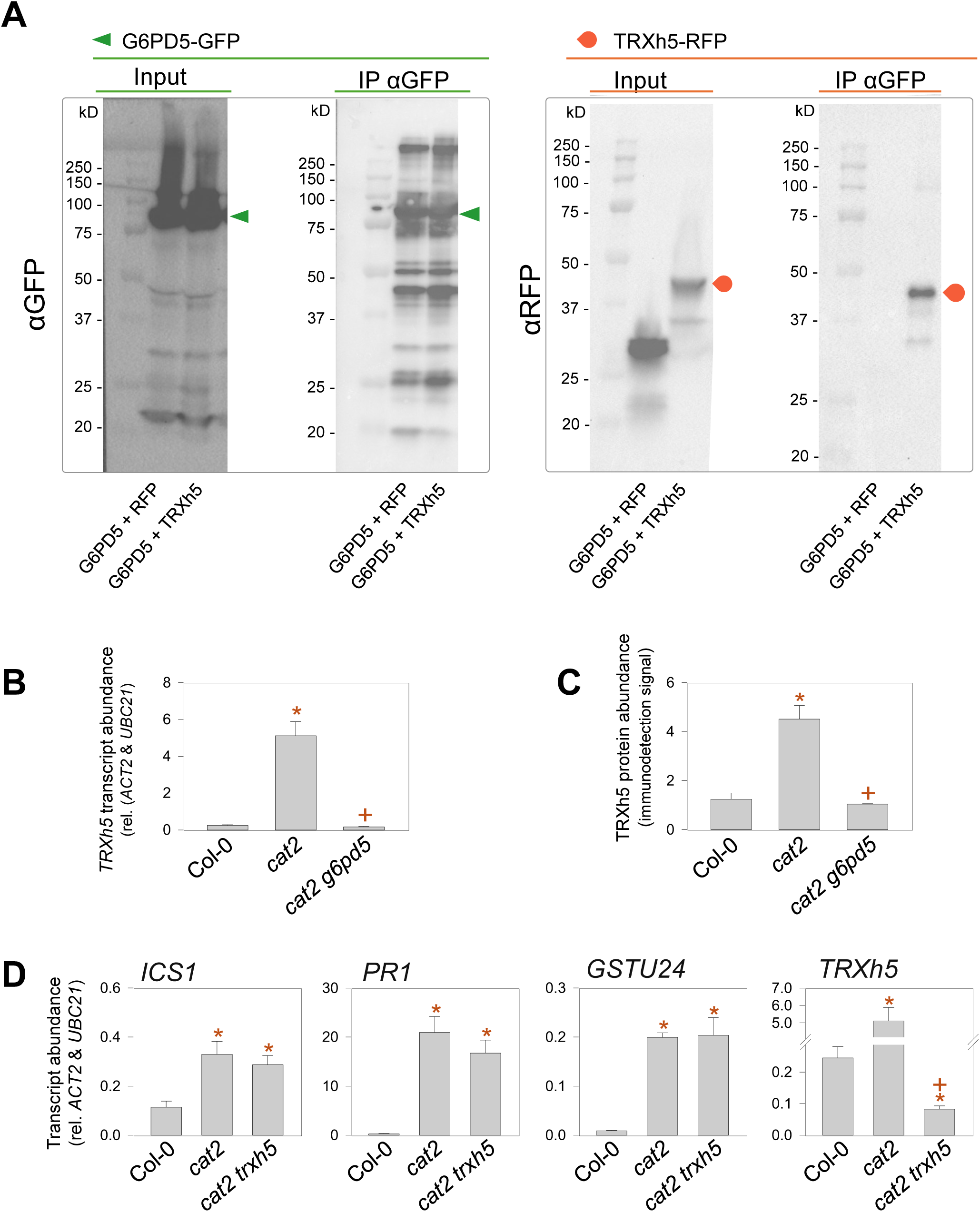
G6PD5 physically associates with TRXh5. A. Western blot against G6PD5-GFP (left) or TRXh5-RFP and RFP alone (right). For the two immunodetections, input refers to the total protein extract (input) and IP G6PD5 to the pulled-down fraction by treatment using αGFP magnetic beads. The different analyses are shown in the same order for the three lanes: molecular weight marker (Marker; kDa: kilodalton), G6PD5-GFP + RFP co-expressed in the plant (G6PD5 + RFP) and G6PD5-GFP + TRXh5-RFP co-expressed in the plant (G6PD5 + TRXh5). The positions of G6PD5-GFP and TRXh5-RFP on the immunoblots are indicated by a green triangle and an orange drop, respectively. B. TRXh5 transcript abundance determined by RT-qPCR analysis. C. TRXh5 protein abundance based on immunodetection specific for the TRXh5 protein using western-blot. The TRXh5 signal is normalized relative to Col-0 signal to allow quantification of relative protein abundance. D. RT-qPCR analysis of *ICS1*, *PR1*, *GSTU24* and *TRXH5* transcript abundance in Col-0, *cat2* and *cat2 trxh5* mutants. For panels B to D, analyses were performed on leaf extracts of three weeks old plants grown in long day. Data are means ± SE of three biological replicates. Asterisks (*) indicate significant differences relative to Col-0, and plus signs (+) indicate significant differences relative to *cat2* (Student’s *t*-test, *p* < 0.05).

TRXh5 catalyzes the reduction and activation of NONEXPRESSOR OF PATHOGENESIS-RELATED GENES 1 (NPR1), a master regulator of SA-dependent *PR* gene expression (Tada et al., 2008). The cytosolic TRXh5-NPR1 module is maintained in a reduced state by NTR, for which the necessary NADPH could be partly or wholly supplied by G6PD5. To investigate the possibility that the role of G6PD5 in the *cat2* context might be simply explained by physical and/or metabolic interaction with the NTR-TRXh5-NPR1 module, we investigated the phenotypes and SA or related factors in *cat2 trxh5* and *cat2 npr1* (Han et al., 2013; Chen et al., 2026) grown in the same conditions as *cat2 g6pd5*. Although *TRXh5* transcript abundance was strongly induced in *cat2*, it was decreased in *cat2 trxh5*, consistent with loss of *TRXh5* expression (Fig. 8D). However, under our growth conditions, loss of *TRXh5* function had only a minor effect on *cat2*-triggered lesion phenotypes (Supplementary Figure S8). Consistent with this, gene expression analysis showed that the SA pathway remained strongly induced in *cat2 trxh5*: in stark contrast to *cat2 g6pd5* (Fig. 2C), *ICS1* and *PR1* transcripts were not significantly different than in *cat2* (Fig. 8D). While the *npr1* mutation did impact SA levels in the *cat2* background, this was in a manner that was opposite to *g6pd5*. Extensive lesions were apparent on *cat2 npr1* leaves, and this effect was associated with very high SA accumulation, to more than three-fold *cat2* values (Supplementary Figure S9).

These observations indicate that, despite the confirmed physical association between G6PD5 and TRXh5, the effect of the *g6pd5* mutation cannot be explained simply by abolition of this function.

## Discussion

As enzymes catalyzing the first steps of the OPPP, G6PDHs are potentially key contributors to processes requiring NADPH. Within the context of oxidative stress, such processes notably include ROS production by NADPH oxidases, ROS removal by the ascorbate-glutathione and other peroxidase-based pathways, and redox signaling via NADPH-requiring thiol-disulfide regulation. As well as these and other NADPH-dependent processes, novel functions for G6PDHs may await discovery. Our aim here was to define the importance of specific G6PDH isoforms in conditions where plants are undergoing oxidative stress due to compromised activity of an important antioxidative enzyme.

### Cytosolic G6PDHs are key determinants of oxidative stress signaling, with a prominent role for G6PD5

Our data establish cytosolic G6PDHs as key determinants of SA signaling triggered by oxidative stress, with G6PD5 emerging as the predominant isoform in this response. Mutation of G6PD5 in the *cat2* background fully abolishes lesions, SA accumulation and associated defense signaling (Figs 1-3). In contrast, the impact of loss of *G6PD6* function is more moderate and produces only partial suppression of *cat2* phenotypes, indicating a more limited contribution of this isoform in conditions of intracellular oxidative stress.

This functional divergence between cytosolic G6PDH isoforms likely reflects differences in catalytic properties, abundance, and tissue-specific expression patterns. Indeed, G6PD6 has a markedly lower affinity for NADP^+^, with a K_m_ of 6500 µM compared to 19 µM for G6PD5 (Wakao and Benning, 2005). Earlier studies reported minimal or no G6PD6 activity in Arabidopsis leaves, consistent with a limited role in leaf metabolism in the absence of stress (Wakao and Benning, 2005). However, several observations from the present study indicate that G6PD6 is expressed and functional to some extent in leaves. First, a significant decrease in extractable leaf G6PDH activity was observed in *g6pd6*. Second, G6PD6 was identified as a potential interactor of G6PD5 in leaf extracts. Third, the loss of *G6PD6* function partially suppresses the *cat2-*phenotype. Together, these observations indicate that G6PD6 is indeed expressed and functional in leaves, and that this isoform makes some contribution to cytosolic G6PDH capacity, although its impact on stress signaling is less pronounced compared to G6PD5.

The influence of the cytosolic G6PD isoforms during oxidative stress stands in marked contrast to the absence of effect of the chloroplastic isoforms. It should be noted that oxidative stress in *cat2* is light-dependent, since it is triggered by decreased capacity to process photorespiratory H_2_O_2_. Unlike the cytosolic isoforms, the chloroplastic G6PDH are down-regulated by disulfide reduction in the light and are thought to be mainly active in the dark (Anderson and Duggan, 1976; Wenderoth et al., 1997; Née et al., 2009). The function of cytosolic G6PDHs and their impact on SA signaling is consistent with previous work identifying the cytosol as a central hub for integrating redox and immune signals (Noctor and Foyer, 2016). Loss-of-function mutations in the cytosolic *GR1*, *DHAR1/2*, and *MDAR2* all attenuate lesion formation, SA accumulation and related defense gene expression in *cat2* (Mhamdi et al., 2010a; Rahantaniaina et al., 2017; Xu et al., 2025). Together with the present work, these studies highlight the importance of cytosolic NADPH generation, glutathione reduction and ascorbate recycling in determining the outcome of oxidative stress.

### The comparison of *g6pd5* and *sid2* effects establishes G6PD5 as a key determinant of oxidative stress-triggered SA signaling

The *sid2* mutation impairs ICS1 activity, which is required for SA accumulation in response to pathogens as well as in *cat2* ((Wildermuth et al., 2001; Chaouch et al., 2010); Fig. 3B). Hence, in *cat2 sid2*, oxidative stress is uncoupled from SA accumulation. The striking similarity between this double mutant and *cat2 g6pd5* suggests that *G6PD5* expression is also required to couple oxidative stress to SA accumulation and other downstream processes. This is evident from the marked overlap between *cat2 g6pd5* and *cat2 sid2* at phenotypic, transcriptional and metabolic levels, where effects associated with the *cat2* mutation largely undergo reversion to the Col-0 state in the double mutants (Fig. 3 to 5).

Effects of oxidative stress in *cat2* on metabolite profiles detected by non-targeted GC-MS profiling have been previously discussed (Noctor et al., 2015). Intriguingly, the metabolite that shows the most dramatic accumulation in *cat2* is gluconate, which may originate from glucose oxidation or by dephosphorylation of 6-phosphogluconate, an indirect product of G6PDH activity, either *in planta* or during GC-MS analysis (Noctor et al., 2015). By contrast, *myo*-inositol shows an opposite effect, with lowest contents in *cat2* (Fig. 4B).This compound is known to be important in cell death regulation (Meng et al., 2009; Donahue et al., 2010) and can suppress lesion formation and SA signaling when supplied exogenously to *cat2* plants (Chaouch and Noctor, 2010). It is notable that like *sid2*, the *g6pd5* mutation partly or wholly reverts *cat2*-triggered effects on these metabolites (Fig. 4B).

While the present study focuses on the role of the G6PDHs in oxidative stress, several previous studies have also linked the OPPP to plant immunity. The *g6pd5 g6pd6* double mutant exhibited reduced induction of defense genes following infection with the nematode *Meloidogyne incognita* (Hu et al., 2019), supporting a broader contribution of cytosolic G6PDHs to defense pathways. In addition, G6PD6 was shown to be phosphorylated by the ASKα kinase in the context of immune signaling (Dal Santo et al., 2012; Stampfl et al., 2016). Although we found no evidence of a significant role for chloroplastic G6PDH isoforms in the present analysis, a knockdown for the plastidial isoform of 6-phosphogluconolactonase (*PGL3*) displayed constitutive induction of SA signaling and enhanced resistance to (hemi)biotrophic pathogens (Xiong et al., 2009). These findings support a model in which factors associated with the OPPP are integrated into immune signaling networks. Our results identify cytosolic G6PD5 as a key component of this network during signaling triggered by intracellular oxidative stress, leading to the question of how this occurs.

### G6PD5 plays a key role in redox homeostasis during oxidative stress

Catalase is unique among H_2_O_2_-processing enzymes because it does not require reductant. Hence, when the major leaf catalase function is lost in *cat2*, a greater load is placed on NADPH-dependent pathways to metabolize H_2_O_2_ processed by CAT2 in the wild-type. Additionally, oxidative stress signaling may involve ROS waves linked to NADPH oxidases (Miller et al., 2010). Based on these concepts, NADPH production by G6PDH could influence oxidative stress responses in two contrasting ways: one that is antioxidant and a second that is pro-oxidant.

While our data reveal a key role for G6PD5 in influencing cell redox state, they provide little evidence that this involves generating reductant for NADPH oxidases. This is because several factors (glutathione redox state, oxidative stress marker transcripts, reactive nitrogen species) indicate that oxidation is increased in *cat2 g6pd5* compared to *cat2* (Fig. 6). Together, these observations indicate that G6PD5-dependent NADPH production may be important in antioxidant processes in the *cat2* context. Indeed, the highly oxidized glutathione status in *cat2 g6pd5* resembles that observed in double mutants in which specific cytosolic NADPH-requiring enzymes of the ascorbate-glutathione pathway have been directly knocked out (Mhamdi et al., 2010a; Xu et al., 2025). This striking effect contrasts with much lesser impact of knocking out genes for other G6PDH isoforms or NADPH-producing enzymes (Mhamdi et al., 2010b; Li et al., 2013; this study); it identifies G6PD5 as a key player in maintenance of redox homeostasis when oxidative load and, therefore, NADPH turnover, are increased. Under such conditions, relief of NADPH inhibition together with increased availability of NADP⁺, a positive regulator of G6PDH activity, may favor OPPP flux (Au et al., 2000; Wakao and Benning, 2005; Tuzet et al., 2019; Wang et al., 2023). Consistent with modeling predictions (Tuzet et al., 2019), our data support the hypothesis that G6PDH activity contributes substantially to NADPH generation during oxidative stress. The functional importance of this aspect of *G6PD5* function in coupling oxidative stress to SA signaling is further studied in the companion manuscript (Trémulot et al.,).

### G6PD5 is integrated into biotic stress and redox protein networks and physically interacts with TRXh5

Our data place G6PD5 at the heart of a network involving several proteins known to be key in redox regulation and SA-related biotic stress responses (Fig. 7; Table 1). The validated interaction with TRXh5 further emphasizes the potential roles of G6PD5 within such networks (Fig. 7 and Fig. 8). TRXh5 is a well-established component of SA signaling. It promotes NPR1 activation and also reduces disulfide bonds and *S*-nitrosylated thiols of other targets (Tada et al., 2008; Kneeshaw et al., 2014). One possibility is that G6PD5 activity is redox-regulated by TRXh5 although current datasets do not identify the presence of Cys switches in the G6PD5 protein. A more likely interpretation is that the association could promote efficient production of NADPH necessary for TRXh5 functions (Laloi et al., 2004; Reichheld et al., 2007). Although reductases such as the cytosolic NTRA were not detected as potential G6PD5 interactors, metabolic coupling between NADPH generation and thiol-based redox systems remains plausible. Besides NPR1 regulation, TRXh5 has also been implicated in controlling GSNOR1 activity and protein *S*-nitrosylation (Zhang et al., 2020; Tabassum and Loake, 2025; Chen et al., 2026) and so G6PD5 might influence GSH/GSNO/NO homeostasis via such pathways. Indeed, loss of G6PD5 function triggers altered glutathione homeostasis and accumulation of RNS (Fig. 6) and GSNOR1 was identified as a potential G6PD5 interactant (Fig. 7).

It should be noted that G6PD5 or TRXh5 could also act independently of their activity; for example, as a scaffold protein or allosteric regulator. A regulatory role independent of the enzyme activity has been reported in other systems (Jin et al., 2022). Notably, human G6PD interacts with enzymes of glutathione synthesis, and our dataset similarly identifies GSH1, the first enzyme of glutathione synthesis, as a potential interactor. Given that Arabidopsis GSH1 is nuclear-encoded and translated in the cytosol prior to plastid import (Wachter et al., 2005), such interaction is mechanistically plausible. This implies an analogous function between kingdoms, which might be related to the importance of G6PD5 in redox homeostasis.

Despite the potential importance of the G6PD5-TRXh5 interaction for redox regulation, our genetic analyses indicate that the interaction alone is not sufficient to explain the strong suppression of SA signaling observed in *cat2 g6pd5*. Most notably, *cat2 trxh5* does not phenocopy *cat2 g6pd5 or cat2 sid2* while the *cat2 npr1* mutants display enhanced, rather than abolished, lesions and SA phenotypes, opposite to the effects observed in *cat2 g6pd5*. Thus, there is little evidence that decreased activity of the TRXh5-NPR1 module is the primary trigger of abolished SA signaling when *G6PD5* expression is compromised.

## Conclusion

Our study reveals a key role for a specific cytosolic G6PDH in determining the outcome of oxidative stress: G6PD5 works to promote cell redox homeostasis, notably countering excessive oxidation of glutathione and interacting with redox regulators such as TRXh5. Alongside these observations, our data show that G6PD5 is required to link oxidative stress to downstream processes such as activation of the SA pathways and defense against biotic stress. Analyses outlined in the accompanying manuscript (Trémulot et al.) provide evidence that the latter occurs through a novel mechanism involving metabolite intermediates.

## Materials and methods

### Plant materials and growth conditions

All *Arabidopsis thaliana* T-DNA lines used in this study were obtained from the Nottingham Arabidopsis Stock Centre (NASC) and were in Columbia-0 (Col-0) ecotype. The analyzed mutant lines were *g6pd1* (GABI_846A05), *g6pd2* (SAIL_1240_G11), *g6pd3* (SALK_139479), *g6pd4* (SALK_131208), *g6pd5* (SAIL_97_F06), *g6pd5-2* (SALK_045083), *g6pd6* (SALK_016157, Wakao and Benning, 2005), *cat2* (SALK_076998, Queval et al., 2007), *sid2* (Wildermuth et al., 2001; Chaouch et al., 2010), *npr1* (Han et al., 2013), *trxh5* (Chen et al., 2026). Double mutants were generated by crossing, and homozygous plants were identified in the F2 generation by genotyping using gene-specific primers as in Supplementary Table S1.

For in-soil growth experiments, seeds were sown on Tref terreau P1 substrate (Jiffy France SARL) and stratified for 3 days at 4°C in the dark. Plants were grown under long days conditions (LD) (16 h light/8h dark) for 3 weeks prior to sampling, unless stated otherwise. Growth chamber conditions were maintained at 20 °C under light, 18 °C in the dark, and 65 % humidity. Light intensity was set to 200 µmol.m^-2^.s^-1^ at leaf level and plants were watered three times per week. All samples were harvested 4 hours after the onset of the light period.

For *in vitro* growth, seeds were sown in 12×12 cm petri dishes containing autoclaved ½ MS and 0.8 % agar medium. The seeds were surface-sterilized by sequential washes with hypochlorite-ethanol (1:9, v/v) and ethanol 95 %, and then stratified for 3 days prior to germination. Plates were placed at a light intensity of 70 µmol.m^-2^.s^-1^ in LD at temperatures of 20 °C (light period) and 18 °C (dark period) and a humidity of 65 %.

Unless stated otherwise, all experiments were performed using three independent biological replicates. Each of the biological repeats was obtained from 3 technical replicates.

### Vector construction and transformation

Total RNA was extracted with ReliaPrep™ RNA Miniprep Systems from leaf tissue (Col-0) to generate cDNA by reverse transcription with qScript™ cDNA Supermix (Quantabio). The coding sequences (CDS) were amplified using gene-specific primers (Supplementary Table S1) and iProof™ High-Fidelity DNA Polymerase (Bio-Rad), with PCR conditions optimized individually for each gene. Sequences were then cloned into the pDONR^TM^221 vector (Invitrogen) using the Gateway^®^ BP recombination reaction, followed by LR recombination reactions using the manufacturer’s instructions (Invitrogen). The vector pK7RWG2 or pK7FWG2 and pH7FWG2 were used, respectively, for transient and stable *Agrobacterium tumefaciens*-mediated plant transformation (Karimi et al., 2002). During cloning, colony PCR was carried out using ALLin™ Red Taq Mastermix (highQu) to confirm the presence and the size of the inserts. All final constructs were verified by Sanger sequencing of the CDS. Plasmids were propagated in E. coli DH5α, grown at 37°C on LB medium supplemented with appropriate antibiotics. Plasmid DNA was isolated using the GeneJET™ Plasmid Miniprep Kit (Thermo Fisher Scientific), following the manufacturer’s instructions.

For the generation of stable transgenic lines, the constructs of interest were introduced into Agrobacterium strain GV3101, and plants were transformed using the floral dip method (Clough and Bent, 1998). T1 seeds were surface-sterilized and selected on growth medium containing the appropriate selection antibiotics. After initial selection, resistant T1 seedlings were transferred to soil and grown to maturity for seed production. Segregation analysis was performed in subsequent generations by germinating surface-sterilized T2 seeds on selective medium. Lines displaying segregation ratios consistent with a single T-DNA insertion were further propagated, and homozygous lines were identified in the following generation using the same selection strategy.

### Transcript quantification by RT-qPCR

Total RNA was extracted using the ReliaPrep™ RNA Tissue Miniprep System (Promega), following the manufacturer’s instructions, with an on-column DNase digestion step included during extraction. RNA concentration was determined using a NanoDrop 2000 spectrophotometer (Thermo Scientific). First-strand cDNA synthesis was performed from 1 µg of total RNA using the qScript™ cDNA Synthesis Kit (Quantabio), according to the manufacturer’s protocol. Prior to quantitative PCR (qPCR), cDNA samples were diluted eight times. The qPCR analysis was performed using LightCycler^®^ SYBR Green I Master mix (Roche) with gene-specific primers (Supplementary Table 1). Transcript abundance was determined using the ΔCt method and using *ACTIN2* (*ACT2*) and *UBIQUITIN-CONJUGATING ENZYME 21* (*UBC21*) as reference genes.

### Transcriptome analysis

Transcriptomic analyses were performed using Affymetrix microarrays. Total RNA extracted from leaf tissue was hybridized to ATH1-121501 GeneChip® arrays by the VIB Nucleomics Core Facility (nucleomicscore.sites.vib.be), following the manufacturer’s instructions (Affymetrix). Raw microarray files (.CEL) were processed using the maEndToEnd workflow (Klaus and Reisenauer, 2018). Quality control was performed using the arrayQualityMetrics 3.64.0 package. Data preprocessing including robust multi-array average (RMA) background correction, quantile normalization, and probe summarization, was performed using the oligo package version 1.71.7 (Carvalho and Irizarry, 2010). Low-intensity probes were filtered using a median log_2_intensity ≥ 4.5 in at least three samples. Probe annotations were retrieved from ath1121501.db using the AnnotationDbi 1.70.0, and only probes with unique TAIR gene mapping were retained for downstream analyses. Differential expression analysis was performed with the limma package (version 3.64.3) by fitting linear models (lmFit) and applying empirical Bayes moderation (eBayes) (Ritchie et al., 2015). For pairwise comparisons, genes were considered differentially expressed when meeting the criteria of a false discovery rate (FDR) < 0.01 and an absolute log₂ fold change > 1 (Supplementary Table S2).

### Targeted quantification of salicylic acid, scopoletin, and camalexin

Quantification of salicylic acid, scopoletin, and camalexin was performed by HPLC, as previously described (Simon et al., 2010). Leaf samples (∼150 mg) were placed in tubes containing metal beads, ground using a Mixer Mill MM400 (Retsch), and kept frozen by repeated addition of liquid nitrogen. Extraction was initiated by adding 1.5 mL of 90 % (v/v) methanol to the ground tissue, followed by centrifugation for 15 min at 16,000 *g* and 4 °C. The supernatant was transferred to a fresh tube, and the pellet was resuspended in 500 µL of 100% methanol and centrifuged again under the same conditions. The resulting supernatant was combined with the first one and centrifuged for 10 min at 16,000 *g* and 4 °C. The final supernatant was transferred to a new tube and evaporated at 30 °C using a Genevac™ miVac concentrator. For analysis of total SA and scopoletin forms, the samples were subjected to acid hydrolysis. Briefly, dried extracts were resuspended in 600 µL of 3 N HCl and incubated at 80 °C for 45 min. After cooling at room temperature, 1 mL of diethyl ether was added, and samples were vortexed for 20 min, followed by centrifugation for 1 min at 16,000 *g* and 4 °C. The upper organic phase was transferred to a new tube. The extraction was repeated once by adding 1 mL of diethyl ether to the remaining aqueous phase. The pooled organic phase was evaporated under a chemical fume hood at room temperature. The dried extract was resuspended in 200 µL of 20 mM sodium acetate buffer (pH 5): acetonitrile (9:1, v/v) and centrifuged for 5 min at 16,000 g prior to HPLC analysis. For fluorescence analysis, 100 µL of each sample was transferred to HPLC vial. For each injection (50 µL), a 25-minute run with an organic-to-aqueous gradient (acetonitrile-sodium acetate buffer) was performed on HPLC (Waters) equipped with NOVA-PAK C18 columns (Waters). Compound contents were quantified based on peak area, with reference to standard calibration curves generated for each experiment.

### Metabolome analyses

Non-targeted gas chromatography-time-of-flight mass spectrometry (GC-TOF-MS) analysis was performed using a method adapted from Noctor et al. (2007). Leaf material (between 100 and 200 mg) was ground to a fine powder in liquid nitrogen. Metabolites were extracted in 80% (v/v) methanol in the presence of two internal standards, α-aminobutyric acid and ribitol. Following centrifugation, multiple 200-µL aliquots of the supernatant were vacuum-dried and stored at -80 °C until analysis. For GC-MS analysis, two successive derivatization steps were performed. The first derivatization consisted of methoximation using 50 µL methoxyamine, followed by silylation using 80 µL N-methyl-N-(trimethylsilyl)trifluoroacetamide (MSTFA). After 2 hours of incubation, 1 µL of the derivatized sample was injected onto the GC column. The total run time per injection was 30 min. Peak identification was performed by comparison with mass spectral databases, and data processing was carried out using LECO Pegasus software. Peak areas were quantified based on selected fragment ions and normalized to the internal standards and sample fresh weight. Statistical analyses were performed using TMEV4 software. Peak areas were mean-centered and scaled by dividing by the standard deviation across all samples prior to hierarchical clustering, and significant metabolites were identified by ANOVA (Supplementary Table S3).

### Redox metabolite profiling

To measure glutathione precursors, a fluorescence-based method was used as described by Queval and Noctor (2007). An aliquot of 200 μL acid extract was mixed with 100 μL 0.2 M CHES buffer (pH 9) and 20 μL of 10 mM DTT to reduce disulfide forms to thiols. After incubation for 30 min at room temperature, 20 µL of 30 mM monobromobimane (MBB) was added to form fluorescent thiol-bimane conjugates. The mixture was incubated for an additional 15 min in the dark, and the reaction was stopped by the addition of 660 µL of 10% (v/v) acetic acid. Samples were centrifuged, filtered, and transferred into HPLC vials, and 50 µL were injected automatically onto the column. Separation was achieved using an elution buffer containing 90 % water, 10 % methanol, and 0.25 % (v/v) HPLC-grade glacial acetic acid. This protocol allowed the separation and quantification of bimane derivatives of Cys, Cys-Gly, and γ-EC. For each compound, standard calibration curves were generated, and data processing was performed using Empower Pro software adapted for the Waters HPLC system.

Glutathione and ascorbate contents and their associated reduction states were quantified according to the detailed protocols described by Noctor and Mhamdi (2022). NAD(H) and NADP(H) pools were quantified as previously described (Mhamdi et al., 2022). All assays were performed in UV-compatible 96-well plates (Costar^®^) using a TECAN Infinite M200 pro plate reader (Tecan Life Sciences).

### Measurements of H_2_O_2_ and reactive nitrogen species

H₂O₂-associated redox changes were monitored using the genetically encoded fluorescent probe roGFP2-Orp1, as previously described (Nietzel et al., 2019; Ugalde et al., 2022). Leaf discs (7 mm) were excised from roGFP2-Orp1-expressing genotypes and their corresponding non-transformed equivalents (Col-0, *cat2* and *cat2 g6pd5*) and immediately transferred to 96-well black plates with clear bottoms containing assay buffer. Leaf discs were incubated in the dark for 2 hours and then transferred to light for 2 hours prior to measurements. Fluorescence was measured using a CLARIOstar® microplate reader (BMG LABTECH). The probe was sequentially excited at 405 nm and 488 nm, and fluorescence emission was recorded at 510-530 nm. Background fluorescence from non-transgenic leaf discs was subtracted prior to analysis. The 405/488 nm fluorescence ratio was calculated for each well and used as a proxy for the oxidation state of roGFP2-Orp1.

The nitric oxide-sensitive fluorescent probe 4,5-diaminofluorescein diacetate (DAF-2DA; Sigma-Aldrich) was used as a cell-permeable reporter of nitric oxide (NO)-derived signals, referred to as reactive nitrogen species (RNS). Leaves were vacuum-infiltrated twice for 15 min at room temperature with 10 µM DAF-2DA solution, then rinsed to remove excess DAF-2DA and incubated in the dark until imaging. Fluorescence emission in the 510-550 nm range was visualized using a confocal laser scanning microscope, with argon laser excitation at 488 nm. Chlorophyll autofluorescence was collected in parallel and used as a control for chloroplast-derived signals.

### G6PDH activity

The G6PDH activity was measured according to Mhamdi et al. (2022). Total protein extracts were prepared in extraction buffer using a mortar and pestle pre-cooled with liquid nitrogen. Protein extracts were desalted using pre-equilibrated NAP-5 columns (Cytiva). Enzymatic activity was assayed by monitoring NADPH production from NADP⁺ as an increase in absorbance at 340 nm, using a Cary60 UV-Vis spectrophotometer (Agilent).

### Protein-protein interaction studies

Immunoprecipitation-mass spectrometry (IP-MS) analysis was performed using *G6PD5-GFP* as bait in the complemented *cat2 g6pd5* background with *cat2* as controls. Detailed protocols were described in Wendrich et al. (2017) and Xu et al. (2025). The *in silico* analysis retained proteins with an enrichment ratio above 50 and a *p*-value for enrichment below 0.01. Gene Ontology (GO) enrichment analysis was performed using the clusterProfiler R package (version 4.16.0) (Wu et al., 2021; Xu et al., 2024). The network visualization was generated in Cytoscape (version 3.10.2) (Shannon et al., 2003), with functional clustering and annotation generated using the AutoAnnotate application (apps.cytoscape.org/apps/autoannotate). Protein functional annotations were retrieved from ThaleMine (Pasha et al., 2020).

Protein-protein interactions were validated for selected candidates by transient co-expression in *Nicotiana benthamiana*, followed by co-immunoprecipitation (Co-IP) targeting GFP-fused G6PD5. Agrobacterium tumefaciens strain GV3101, transformed with G6PD5–GFP, RFP, or TRXh5-RFP constructs, was grown in YEB medium supplemented with appropriate antibiotics at 30 °C until reaching the mid-log phase. Bacterial cultures were pelleted by centrifugation for 30 min at 4,000 *g* and resuspended to an OD_600_ of 1.5 in infiltration buffer containing 100 mM MES (pH 5.6), 10 mM MgCl₂, and 1 µM acetosyringone. Bacterial suspensions corresponding to each construct were co-infiltrated into *N. benthamiana* leaves, together with a P19 silencing suppressor strain, at a 1:1:1 (v/v) ratio, and tissues were harvested 3 days post-infiltration. Proteins were extracted from frozen tissue using liquid nitrogen and an extraction buffer containing 150 mM Tris-HCl (pH 7.5), 150 mM NaCl, 10 mM EDTA, 1 mM sodium molybdate, 1% (v/v) NP-40, 10% (v/v) glycerol, and a protease inhibitor cocktail (Roche). Extracts were clarified by centrifugation for 1 hour at 17,000 *g* and 4°C. For Co-IP, 1 mL of protein extract was incubated for 3 hours at 4 °C with 30 µL of equilibrated GFP-Trap magnetic beads (Chromotek). Beads were collected using a magnetic rack and washed three times with 20 mM Tris-HCl (pH 7.5), 150 mM NaCl, and 0.5% (v/v) NP-40. Bound proteins were eluted by boiling for 10 min at 95 °C. Input and immunoprecipitated fractions were analyzed by immunoblotting using an HRP-conjugated anti-GFP antibody (Miltenyi Biotec) or an anti-RFP antibody (Chromotek) followed by an HRP-conjugated anti-mouse secondary antibody. Proteins were separated on Mini-PROTEAN® TGX™ Stain-Free precast gels and transferred to PVDF membranes (Bio-Rad). Detection was performed using the Western Lightning chemiluminescence kit (GE Healthcare), and signals were visualized with a Bio-Rad ChemiDoc imaging system.

### Pathogen Tests

The virulent bacterial pathogen *Pseudomonas syringae* pv. *tomato* strain DC3000 was used for disease resistance assays at a final inoculum concentration of 5 × 10⁵ colony-forming units (cfu) mL⁻¹. Whole leaves were infiltrated using a 1-mL needleless syringe. Leaf discs (0.5 cm²) were harvested 24 and 48 hours post-inoculation from infiltrated leaves. Four biological samples were generated by pooling two leaf discs each from a different treated plant. Bacterial growth was determined by homogenizing leaf discs in 400 µL of sterile water, followed by plating appropriate serial dilutions on King’s B agar medium supplemented with rifampicin and kanamycin. Colony-forming units were counted after 3 days of incubation, and bacterial titers were calculated accordingly.

### Bioinformatics and statistical analyses

R 4.5.1 in RStudio was used for statistics. Heatmap construction was done with the ComplexHeatmap 2.24.1 package (Gu, 2022), and Venn diagrams were generated using the eulerr 7.0.2 package. Enrichment analysis was performed using clusterProfiler 4.16.0 (Wu et al., 2021), with genes with transcripts sequenced in our microarray experiment as background and a *p*-value cut-off of 0.05 for the hypergeometric over-representation test. Concerning annotation, the AnnotationHub 3.16.1 and org.At.tair.db 3.21.0 packages were used. Plots were generated using the enrichplot 1.28.4 package, ggplot2 and Excel (Microsoft).

### Accessions

The *Arabidopsis thaliana* genes analyzed in this study correspond to the following AGI locus identifiers: *G6PD1* (AT5G35790), *G6PD2* (AT5G13110), *G6PD3* (AT1G24280), *G6PD4* (AT1G09420), *G6PD5* (AT3G27300), *G6PD6* (AT5G40760), *CAT2* (AT4G35090), *ISOCHORISMATE SYNTHASE 1* (*ICS1*; AT1G74710), *PATHOGENESIS-RELATED 1* (*PR1*; AT2G14610), *THIOREDOXIN h5* (*TRXh5*; AT1G45145), and *NONEXPRESSOR OF PR GENES 1* (*NPR1*; AT1G64280).

## Acknowledgments

The authors thank Cécile Raynaud and Catherine Bergounioux for their advice on the cloning procedures. We are grateful to Andreas J. Meyer and Markus Schwarzländer for the generous gift of the roGFP2-Orp1 plasmids.

## Author Contributions

A. M. and G.N. designed the research. L.T and A.M. performed the experiments with contribution from E.I-B. and B.D.R. B.D.R, K.V.D.K., E.I-B and J-P.R. contributed to data analysis and interpretation. G.N. supervised the work with contributions from F.V.B. and A.M. L.T., A.M. and G.N. wrote the manuscript. All authors have read and approved the final manuscript.

## Funding

Lug Trémulot was supported by a PhD fellowship from the Université Paris-Saclay and Ministère de l’Enseignement Supérieur et de la Recherche, France. The GN laboratory received financial support from the French Agence Nationale de la Recherche HIPATH project (ANR-17-CE20-0025) and the Institut Universitaire de France (IUF). The FVB laboratory is supported in part by the Research Foundation Flanders (FWO) The FVB laboratory is supported in part by the Research Foundation Flanders (FWO) (The Excellence of Science [EOS] Research project 869 30829584), and NUCLEOX (grant number G007723N).

## Data availability

The data that support the findings of this study are available from the corresponding author upon reasonable request.

## Supplementary data

The following materials are available in the online version of this article.

**Supplementary Figure S1.**
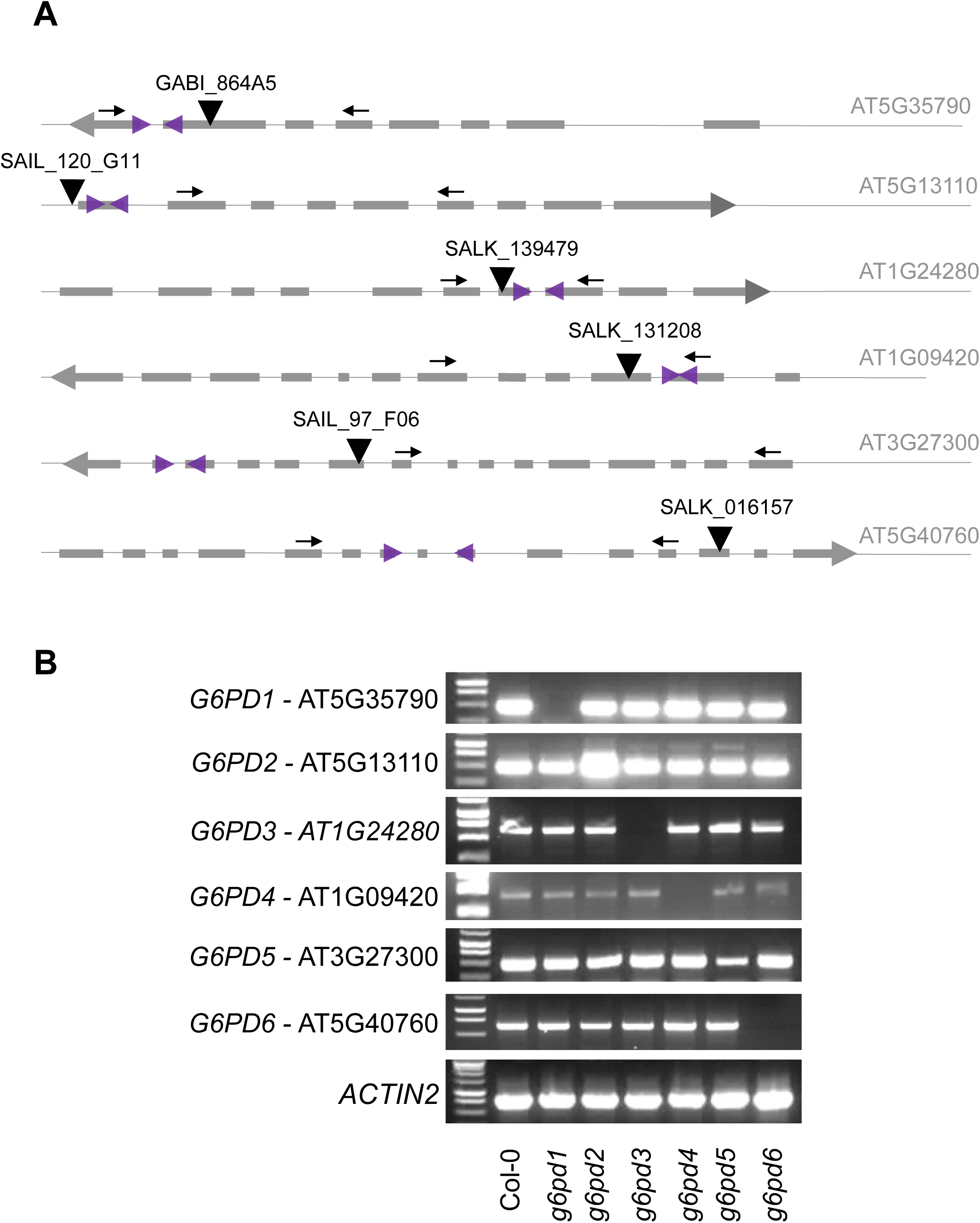
Characterization of *g6pd* mutants. A. Schematic representation of gene structures and primer positions. Exons are shown as grey boxes and introns as connecting lines. Arrows indicate gene orientation, showing transcription in either the forward or reverse direction. Black triangles represent T-DNA insertion sites in the different mutant alleles. Black arrows indicate the positions of primers used for RT-PCR analysis. Purple triangles sitting on the exons indicate the positions of primers used for RT-qPCR analysis. B. RT-PCR analysis of *G6PD* transcripts in Col-0 and *g6pd* mutants. Transcript levels were assessed using the primer pairs indicated in (A). Total RNA was extracted from leaves, reverse-transcribed, and amplified using gene-specific primers. Each row corresponds to a different primer pair targeting a specific transcript. The presence or absence of amplification bands reflects transcript accumulation and integrity in the different genotypes.

**Supplementary Figure S2.**
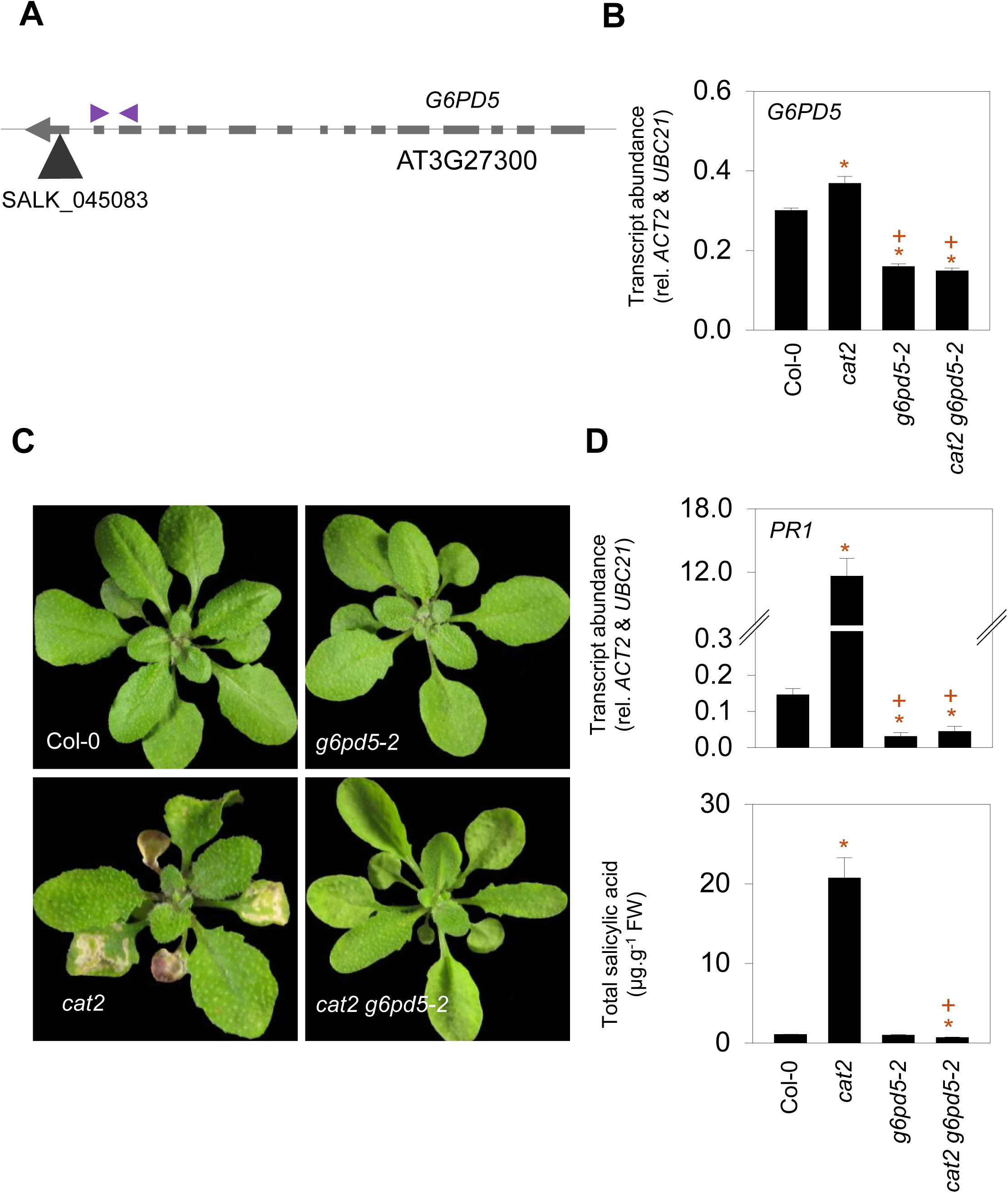
Analysis of *g6pd5* allelic lines. A. Gene structure and primer positions. Schematic representation of *G6PD5* showing exons (grey boxes) and introns (lines). The arrow indicates gene orientation. The black triangle marks the T-DNA insertion site of the allelic line (*g6pd5-2*). Purple arrows indicate the positions of primers used for RT-qPCR analysis. B. *G6PD5* transcript abundance quantified by RT-qPCR using the primer pairs indicated in (A). Transcript abundance is expressed relative to the reference genes *ACT2* and *UBC21*. C. Photographs of plants growing in LD for three weeks. B. Quantification of *PR1* transcripts and total salicylic acid in the different genotypes. In B, and D data are means ± SE of three biological replicates. Asterisks (*) indicate significant differences relative to Col-0, and plus signs (+) indicate significant differences relative to *cat2* (Student’s *t*-test, *p* < 0.05).

**Supplementary Figure S3.**
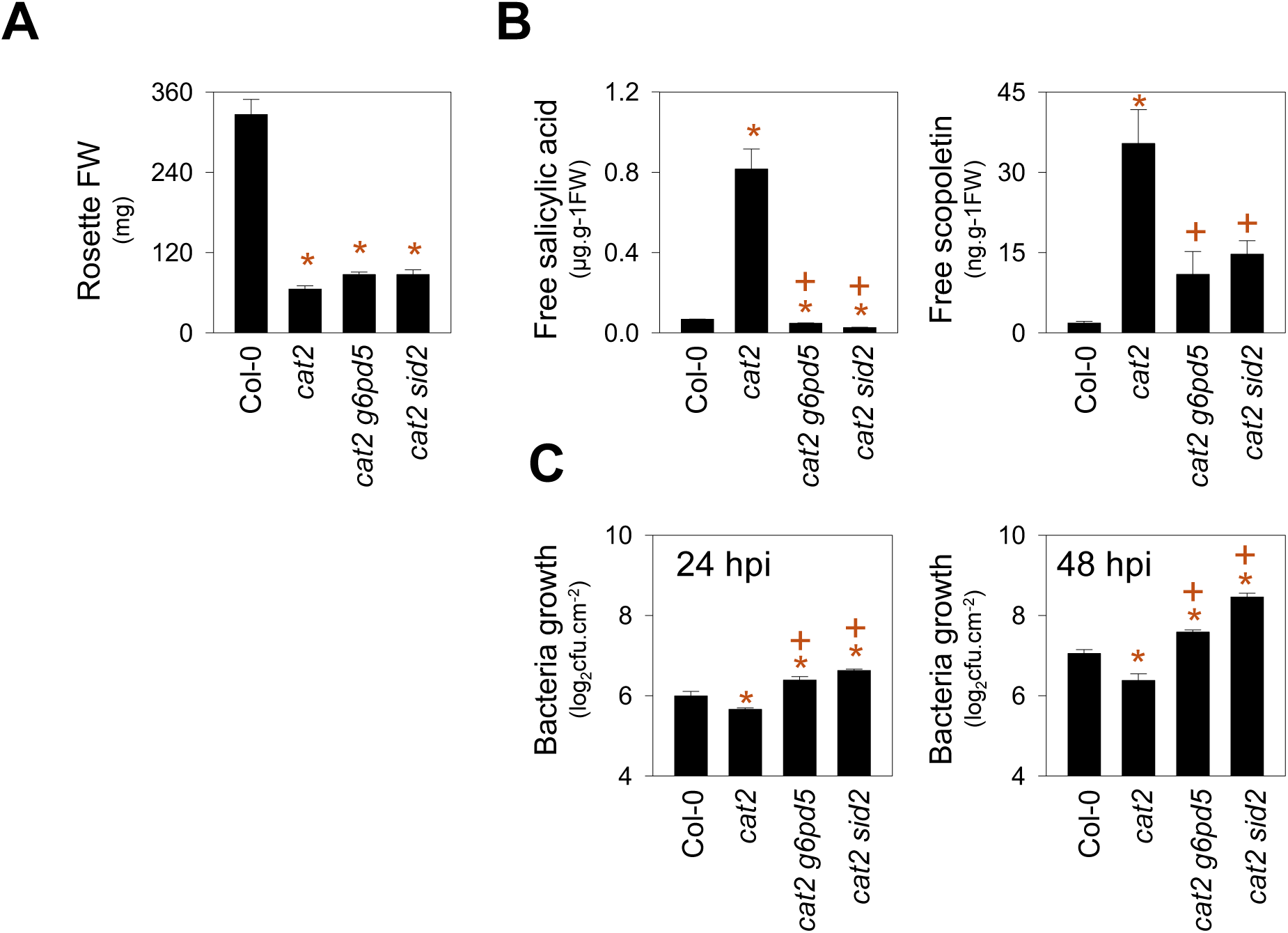
Comparison of *cat2 g6pd5* and *cat2 sid2* mutants. A. Rosette fresh weight. B. Quantification of free salicylic acid and free scopoletin. C. Growth of *Pst* DC3000 at 24 and 48 hpi (hours post-inoculation). Asterisks (*) indicate significant differences relative to Col-0, and plus signs (+) indicate significant differences relative to *cat2* (Student’s *t*-test, *p* < 0.05).

**Supplementary Figure S4.**
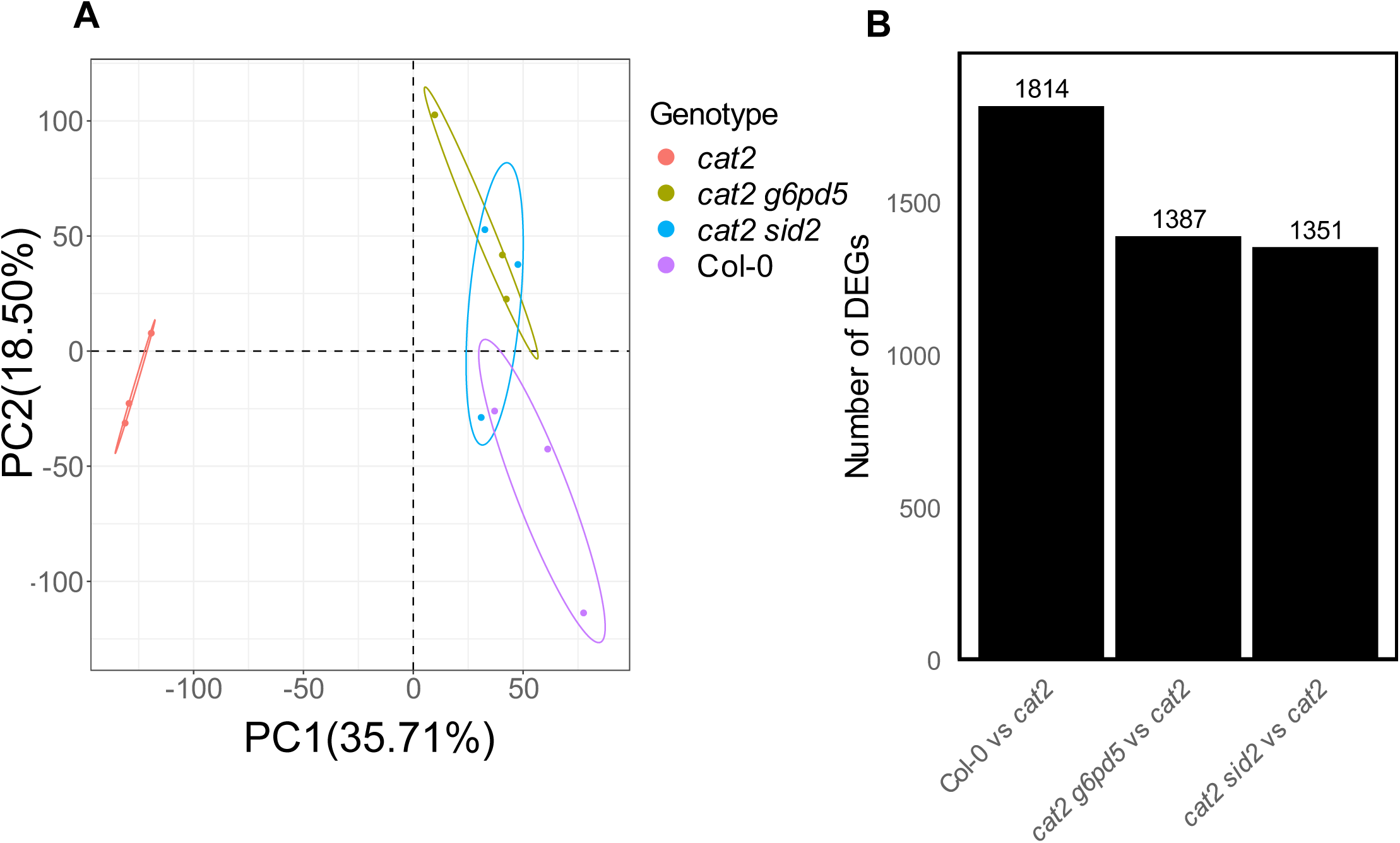
Comparison of transcriptome profiles. A. Principal component (PC) analysis of transcriptome datasets showing the distribution of biological replicates in the four genotypes, Col-0, *cat2*, *cat2 g6pd5*, and *cat2 sid2*. Each point represents an individual biological replicate. Ellipses indicate the distribution of replicates for each genotype. The analysis highlights the separation of *cat2* from the other genotypes and the similarity between *cat2 g6pd5* and *cat2 sid2* relative to *cat2*. B. Number of differentially expressed genes (DEGs) identified in pairwise comparisons relative to *cat2* (Col-0 vs *cat2*, *cat2 g6pd5* vs *cat2*, and *cat2 sid2* vs *cat2*).

**Supplementary Figure S5.**
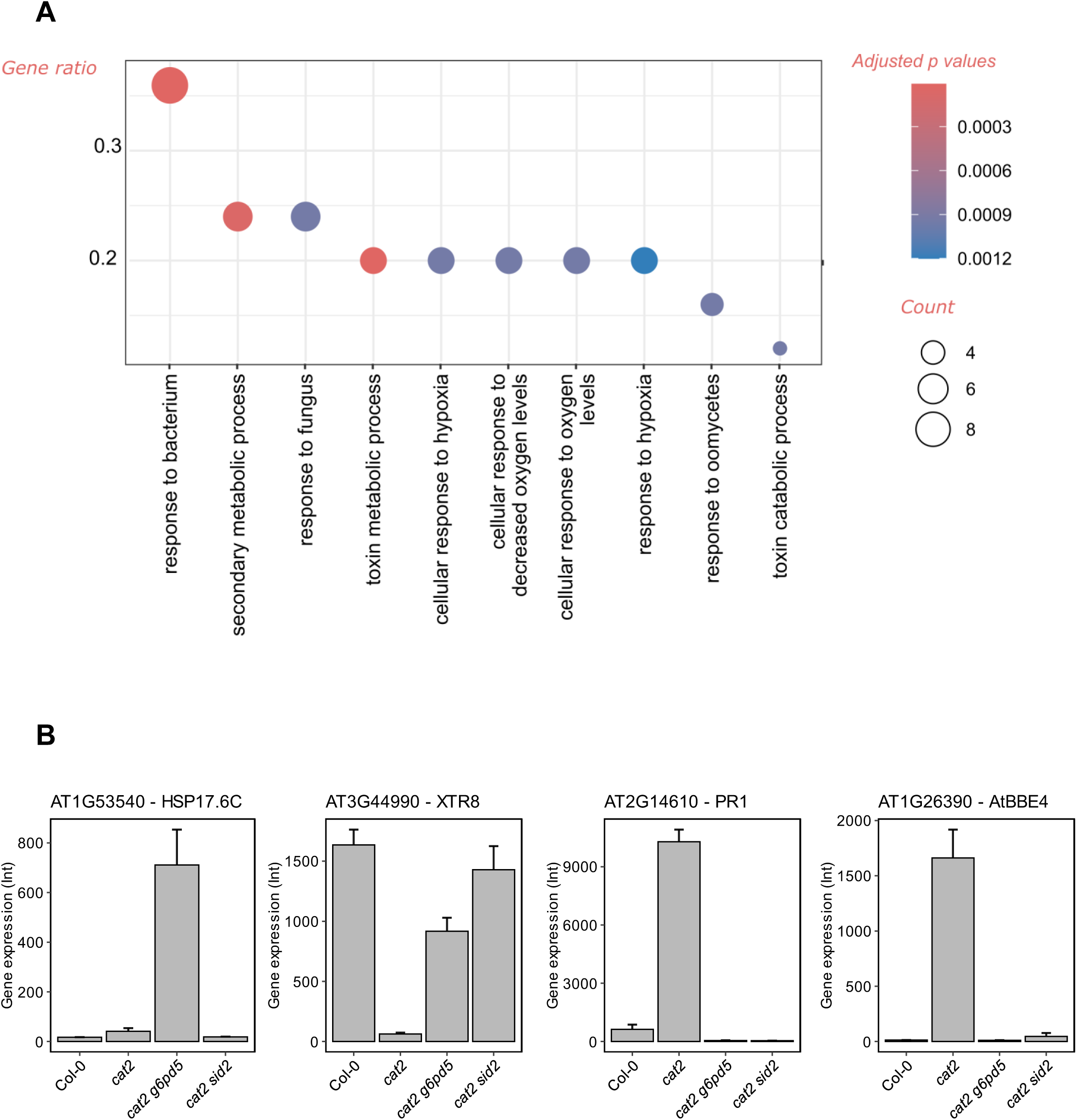
Transcriptomic signatures associated with impaired SA signaling in *cat2 g6pd5*. A. Gene Ontology (GO) enrichment analysis of differentially expressed genes identified in the *cat2 g6pd5* vs *cat2* comparison. Only top 10 biological processes are shown (*p*-adjust < 0.05). Bubble size indicates the number of genes contributing to the enrichment, and color represents adjusted *p*-values. GeneRatio corresponds to the proportion of DEGs associated with a given GO term relative to the total number of genes annotated for that GO term. B. Expression profiles of the two genes showing the highest and lowest fold changes in the *cat2 g6pd5* versus *cat2* comparison. Gene expression values (intensity, Int) are derived from microarray analysis

**Supplementary Figure S6.**
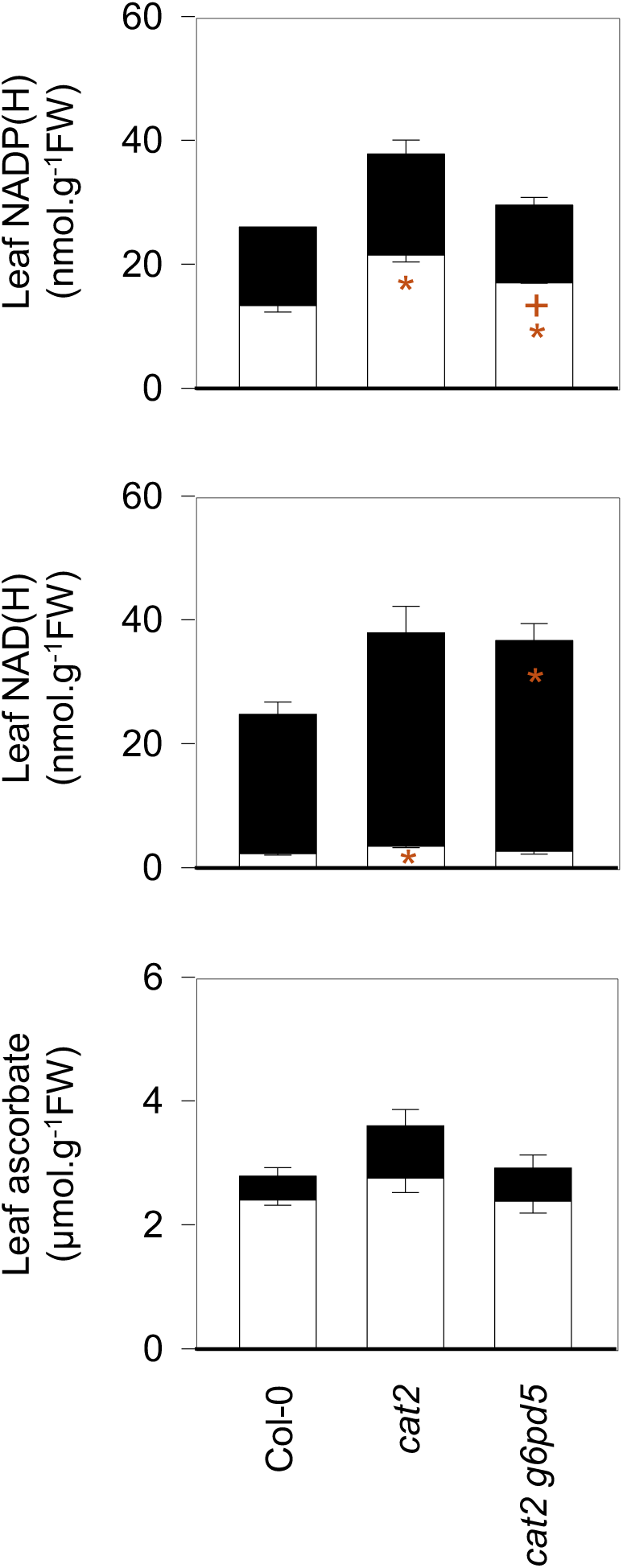
Effects of G6PD5 impairment on ascorbate, NAD(H) and NADP(H) levels in Col-0, *cat2* and *cat2 g6pd5*. White bars indicate reduced forms and black bars indicate oxidized forms of each metabolite. Data are mean ± SE of three biological replicates. Asterisks (*) indicate significant differences relative to Col-0, and plus signs (+) indicate significant differences relative to *cat2* (Student’s *t*-test, *p* < 0.05).

**Supplementary Figure S7.**
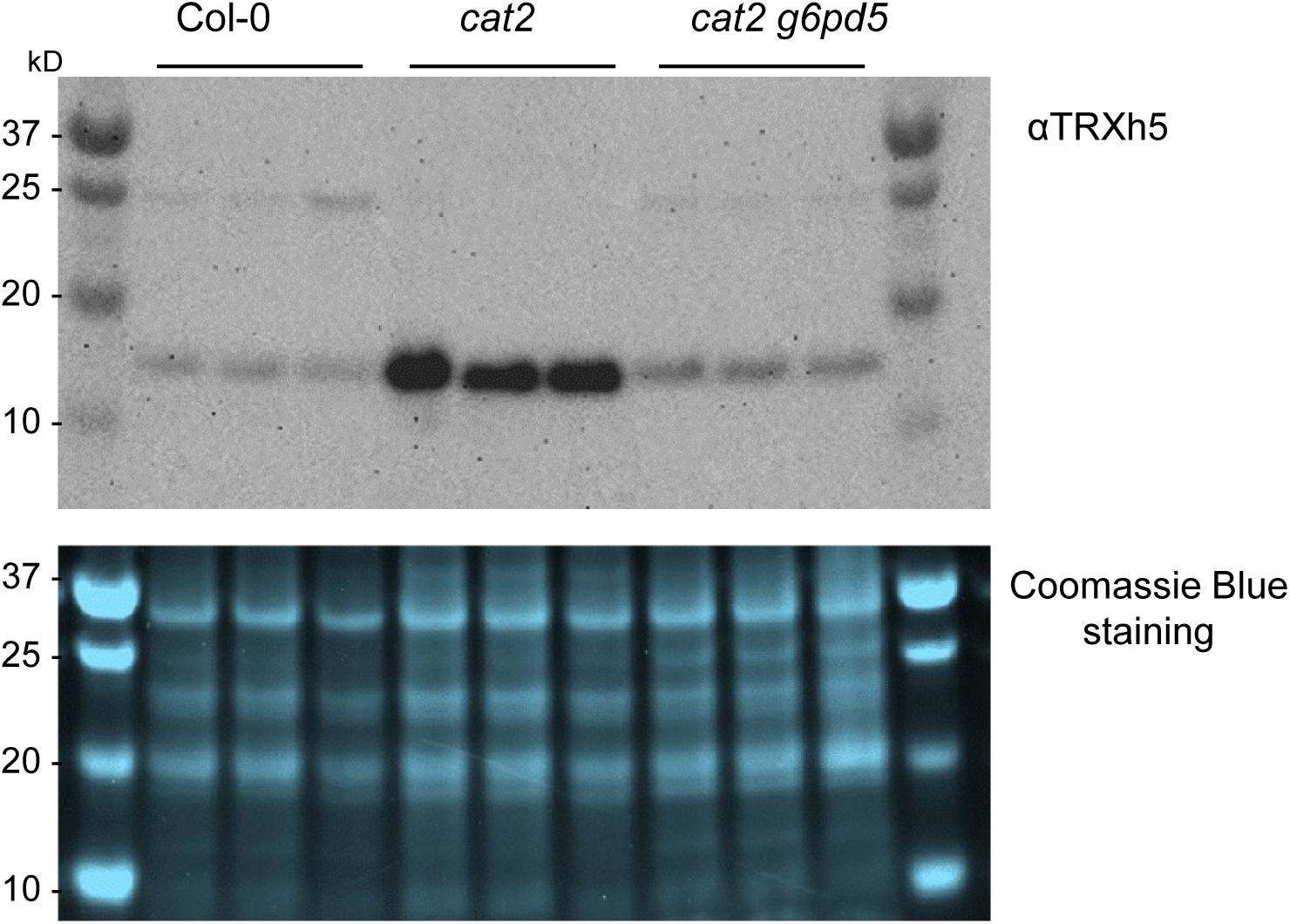
Immunoblot analysis of TRXh5 protein abundance. Western blot analysis of TRXh5 protein abundance in the indicated genotypes (upper panel). Total protein extracts were separated by SDS-PAGE and probed with an anti-TRXh5 antibody. The predominant band detected corresponds to the expected molecular mass of TRXh5 (∼13.2 kDa). Molecular weight markers are shown at the left and right of the blot. Coomassie Blue staining (lower panel) of the corresponding SDS-PAGE gel showing comparable protein loading across samples.

**Supplementary Figure S8.**
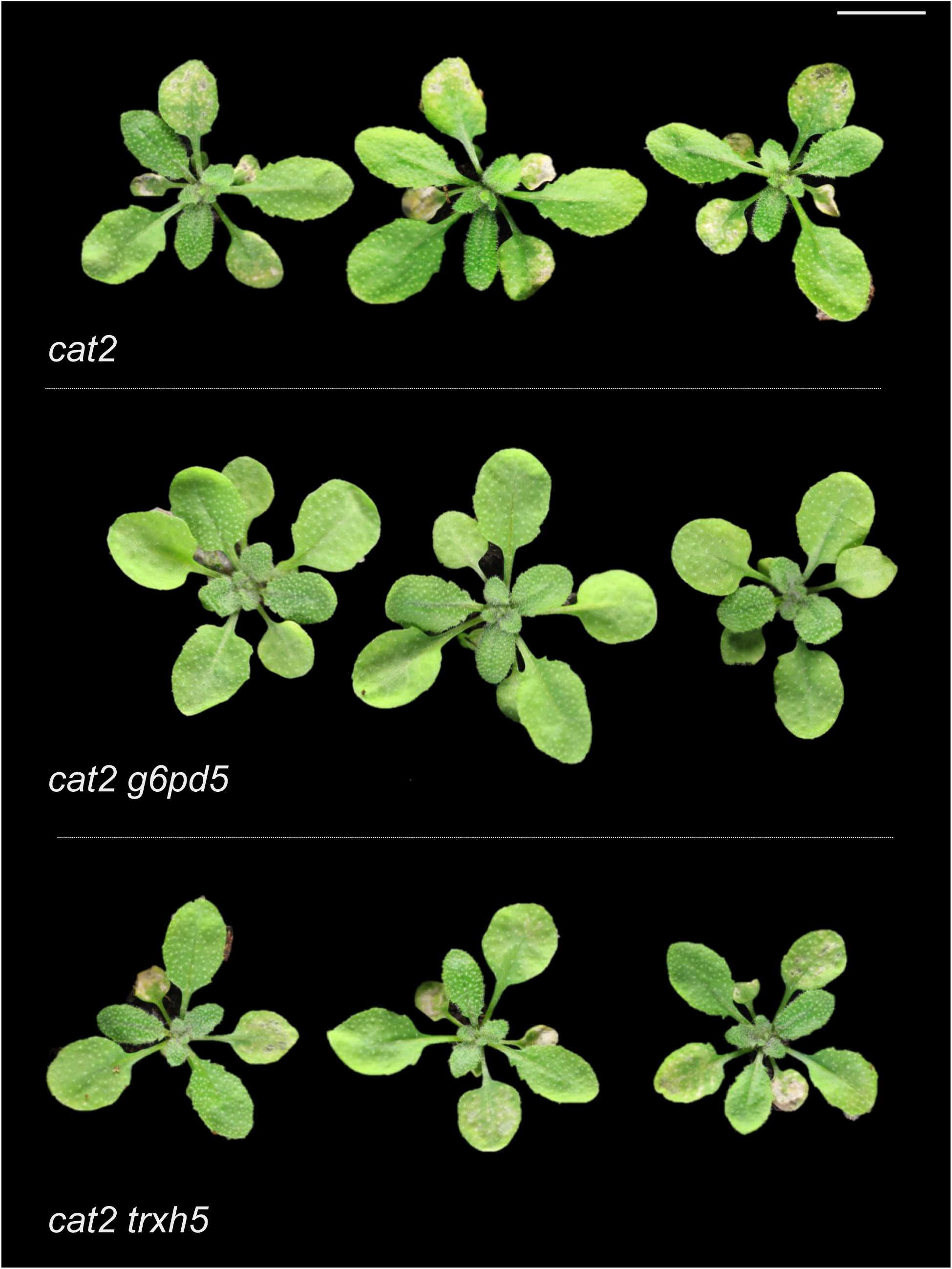
Impact of *TRXh5* mutation on *cat2-*triggered lesion formation. A. Representative photographs of Col-0, *cat2*, and *cat2 trxh5*. The *cat2* mutant displays characteristic lesion phenotype that is maintained in the *cat2 trxh5* background. Scale bars = 2 cm.

**Supplementary Figure S9.**
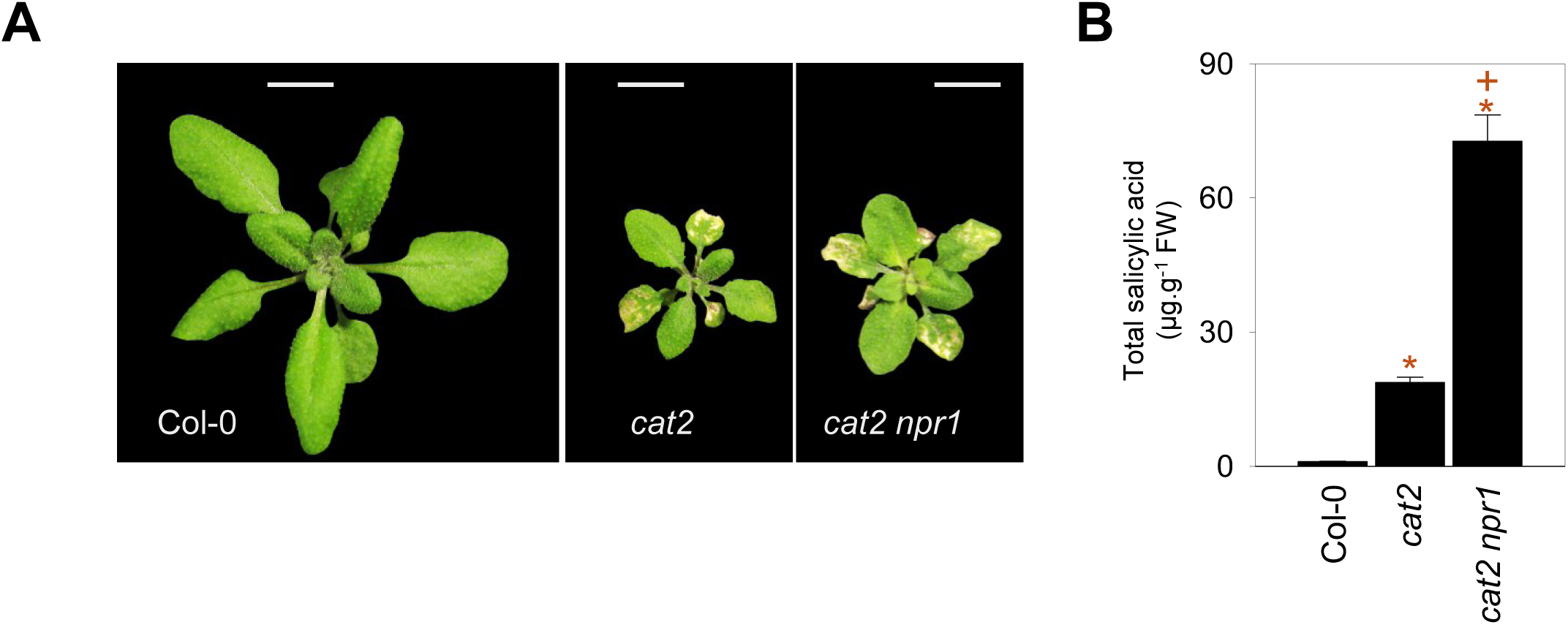
Impact of *NPR1* mutation on *cat2-*triggered lesion formation and SA accumulation. A. Representative photographs of Col-0, *cat2*, and *cat2 npr1*. The *cat2* mutant displays characteristic lesion formation, a phenotype that is enhanced in the *cat2 npr1* background. Scale bars = 2 cm. B. Quantification of total SA levels in the indicated genotypes. Data represent mean ± SE of three biological replicates. Asterisks (*) indicate significant differences relative to Col-0, and plus signs (+) indicate significant differences relative to *cat2* (Student’s *t*-test, *p* < 0.05).

## Supplementary tables

**Supplementary Table S1.** List of primers used in this study.

**Supplementary Table S2.** Differentially expressed metabolites in all genotypes.

**Supplementary Table S3.** Differentially expressed genes in all genotypes.

**Supplementary Table S3.** List of identified potential interactors.

## Notes

### Competing Interest Statement

The authors have declared no competing interest.

